# Conserved loop of a phase modifier endows protein condensates with fluidity

**DOI:** 10.1101/2024.07.03.601791

**Authors:** Honoka Kawamukai, Motonori Matsusaki, Takanari Tanimoto, Mai Watabe, Ken Morishima, Shunsuke Tomita, Yoichi Shinkai, Tatsuya Niwa, Taro Mannen, Hiroyuki Kumeta, Hitoki Nanaura, Kotona Kato, Takuya Mabuchi, Yuichiro Aiba, Takeru Uehara, Noriyoshi Isozumi, Yoshika Hara, Shingo Kanemura, Hiroyoshi Matsumura, Kazuma Sugie, Koichiro Ishimori, Takahiro Muraoka, Masaaki Sugiyama, Masaki Okumura, Eiichiro Mori, Takuya Yoshizawa, Tomohide Saio

**Affiliations:** Graduate School of Chemical Sciences and Engineering, Hokkaido University, Sapporo, Hokkaido 060-8628, Japan; Graduate School of Medical Sciences, Tokushima University, Tokushima, Tokushima 770-0042, Japan; Institute of Advanced Medical Sciences, Tokushima University, Tokushima, Tokushima 770-0042, Japan; College of Life Sciences, Ritsumeikan University, Kusatsu, Shiga 525-8577, Japan; Frontier Research Institute for Interdisciplinary Sciences, Tohoku University, Sendai, Miyagi 980-8578, Japan; Institute for Integrated Radiation and Nuclear Science, Kyoto University, Sennan-gun, Osaka 590-0494, Japan; Health and Medical Research Institute, National Institute of Advanced Industrial Science and Technology (AIST), Tsukuba, Ibaraki 305-8566, Japan; Molecular Neurobiology Research Group, Biomedical Research Institute, National Institute of Advanced Industrial Science and Technology (AIST), Tsukuba, Ibaraki 305-8566, Japan; Cell Biology Center, Institute of Innovative Research, Tokyo Institute of Technology, Yokohama, Kanagawa 226-8501, Japan; Global Station for Soft Matter, Global Institution for Collaborative Research and Education, Hokkaido University, Sapporo, Hokkaido 011-0021, Japan; Department of Neurology, Nara Medical University, Kashihara, Nara 634-8521, Japan; Department of Future Basic Medicine, Nara Medical University, Kashihara, Nara 634-8521, Japan; Faculty of Medicine, Tokushima University, Tokushima, Tokushima 770-0042, Japan; Institute of Fluid Science, Tohoku University, Sendai, Miyagi 980-8577, Japan; Department of Chemistry, Graduate School of Science, Nagoya University, Nagoya, Aichi 464-8602, Aichi, Japan; Department of Applied Chemistry, Graduate School of Engineering, Tokyo University of Agriculture and Technology, Koganei, Tokyo 184-8588, Japan; Department of Chemistry, Faculty of Science, Hokkaido University, Sapporo, Hokkaido 060-0810, Japan; Institute of Global Innovation Research, Tokyo University of Agriculture and Technology, Fuchu, Tokyo 183-8538, Japan; Kanagawa Institute of Industrial Science and Technology (KISTEC), Ebina, Kanagawa 243-0435, Japan; V-iCliniX Laboratory, Nara Medical University, Kashihara, Nara 634-8521, Japan

**Author notes:** These authors contributed equally (Honoka Kawamukai, Motonori Matsusaki). To whom correspondence should be addressed: Tomohide Saio: Institute of Advanced Medical Sciences, Tokushima University, Tokushima, Tokushima 770-0042, Japan; Takuya Yoshizawa: College of Life Sciences, Ritsumeikan University, Kusatsu, Shiga 525-8577, Japan; Eiichiro Mori: Department of Future Basic Medicine, Nara Medical University, Kashihara, Nara 634-8521, Japan. Research Division, Chugai Pharmaceutical, Yokohama, Kanagawa, 244-8602, Japan (Takuya Yoshizawa).

## Abstract

Dipeptide repeats (DPRs) that are gene products from abnormal hexanucleotide repeat expansion in *C9orf72* trigger amyotrophic lateral sclerosis (ALS) through unknown mechanism. This study highlights, importin Karyopherinβ2 (Kapβ2), which is responsible for nuclear transport and phase modification of RNA-binding proteins (RBPs), as a major DPR target. We demonstrate DPR accumulation in the nucleus via Kapβ2-mediated transport, which results in dose-dependent toxicity observed in nematode and yeast models. In vitro interaction studies exploiting chemical probe arrays and biophysical measurements reveal multivalent DPR binding to Kapβ2, including at the conserved acidic loop. Refractive index and fluorescence imaging coupled with biochemical assays unveiled that binding of excess DPRs to the acidic loop turns a phase modifier Kapβ2 into phase disrupter, resulting more condensed and viscous RBP condensates. Our findings provides molecular insight into *C9orf72*-ALS related to age and repeat expansion.

---

The discovery of the association between *C9orf72* gene and familial amyotrophic lateral sclerosis (ALS) and familial frontotemporal dementia (FTD) marked a pivotal advancement in neurodegenerative research^1,2^. Patients with ALS and FTD show 700– 1600 repeats of an abnormal extension of the hexanucleotide GGGGCC in intron 1 of *C9orf72* compared to healthy individuals^1^. This expanded repeat sequence undergoes repeat-associated non-ATG initiated (RAN) translation^3,4^, generating various di-peptide repeats (DPRs)^5^. Of these DPRs, the Arg-containing glycine–arginine (GR) and proline– arginine (PR) exhibit neurotoxicity from various action points, including inhibition of protein translation and nuclear transport^6–12^. Despite extensive research, the detailed mechanisms underlying DPR-induced neurotoxicity remain elusive. As possible mechanisms, recent studies have focused on dysregulation of nuclear transport and liquid-liquid phase-separation (LLPS), both of which involve the importin Karyopherinβ2 (Kapβ2; also known as Transportin 1)^13^. In nuclear transport, Kapβ2 recognizes the proline-tyrosine nuclear localization signal (PY-NLS) of cargo proteins and transports them into the nucleus^14,15^. This is followed by cargo release in the nucleus upon binding of the low molecular weight G protein Ran (RanGTP)^16,17^. In LLPS regulation, Kapβ2 works as a phase modifier through selective and multivalent recognitions of PY-NLS-containing RNA-binding proteins (RBPs), including fused in sarcoma (FUS) and hnRNPA2, to resolve LLPS droplets^18^. The multivalent recognition of RBPs by Kapβ2 includes recognition of PY-NLS, low-complexity (LC) region, arginine-glycine-glycine (RGG) domain, RRM domain, and ZnF domain^13,18^. A recent study revealed that DPRs bind to the NLS-binding site of Kapβ2 and inhibit its function as a phase modifier^19^. However, the molecular mechanism for dose-dependent and length-dependent toxicity of DPRs in *C9orf72*-related ALS pathogenesis remains to be elucidated. Given the DPR accumulation in the nucleus to inhibit RNA metabolism and protein expression^20,21^, the mechanisms of DPR accumulation in the nucleus should also be the key.

In this study, we aimed to elucidate the dose-dependent effect of DPRs on Kapβ2 activity and structure. Through an integration of in vivo assays using a nematode ALS model and yeast cells with in vitro assays using a chemical sensor array “chemical tongue”, biophysical measurements, refractive index (RI) imaging, and structural investigations utilizing nuclear magnetic resonance (NMR), we demonstrate the multivalent binding of excess DPRs to Kapβ2, including the interaction with the conserved acidic loop. Importantly, our findings highlight the disruptive impact of DPR binding on Kapβ2’s physical properties, leading to its accumulation and sequestration in RBP condensates. This accumulation in turn impedes the nuclear import of RBPs, reminiscent of the pathological hallmarks observed in ALS neurons^22–24^. Our study sheds light on the intricate interplay between DPRs and Kapβ2, providing mechanistic insights into *C9orf72*-related ALS pathogenesis and offering potential therapeutic targets.

## PR poly-dipeptides ride nuclear import receptor Kapβ2 to accumulate in the nucleus and express toxicity

We first aimed to investigate if the nuclear import receptor Kapβ2 is involved in the nuclear accumulation of DPRs. GFP-PR20-HA was localized in the nucleus when expressed in HeLa cells, whereas co-expression of Kapβ2-specific peptide inhibitor M9M^25^ decreased PR20 transportation into the nucleus (Fig. 1A; Extended Data Fig. 1), indicating that the nuclear transport of PR20 is partially mediated by Kapβ2. Consistent with this, an in vitro cargo-unloading experiment showed that maltose-binding protein (MBP)-PR18 bound to Kapβ2 was unloaded and replaced by RanGTP, as is the case with the physiological cargo hnRNPA2 NLS (Fig. 1B, Extended Data Fig. 2). These data suggest that DPRs rides on the Kapβ2 nuclear import system to be accumulated in the nucleus. The nuclear accumulation of PR20 but not PR8 was also observed in yeast cells (Extended Data Fig. 3A). Toxicity of PR poly-dipeptide expression on yeast growth was observed with PR20 but not with PR8, indicating length-dependent toxicity (Extended Data Fig. 3B). Moreover, yeast growth curves with different PR20 expression levels showed that the higher the expression level, the slower the growth, demonstrating the dose-dependent toxicity of PR20 (Fig. 1C, Extended Data Fig. 3C).

**Figure 1.**
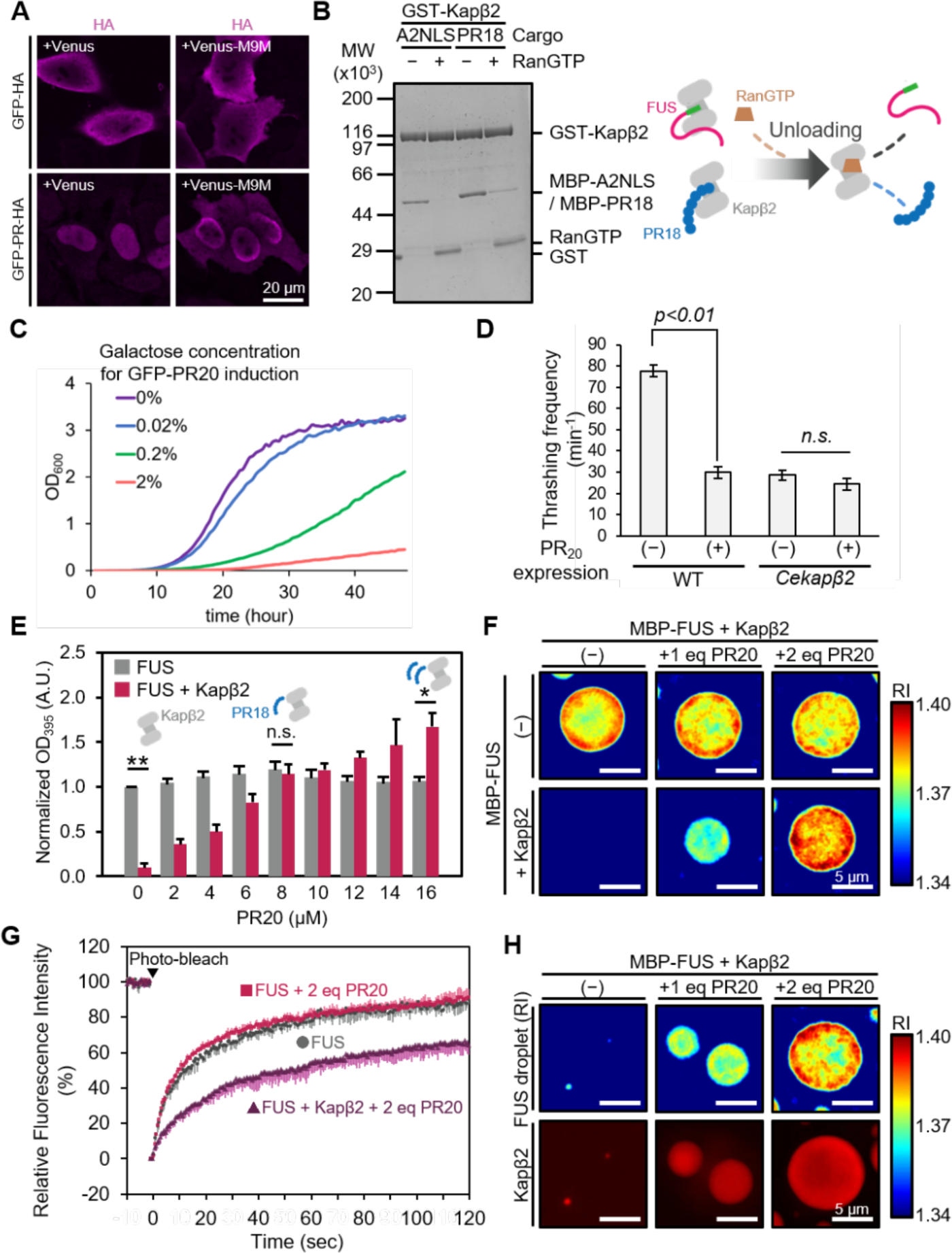
The excess PR poly-dipeptides are transported to the nucleus as Kapβ2 cargo and exhibit toxic effect in vivo. **A**. Subcellular localization of the PR20 peptide under transportin inhibition by M9M peptide. **B**. Pull-down binding assay of MBP-hnRNPA2NLS or MBP-PR18 with immobilized GST-Kapβ2 in the presence of RanGTP. Right panel shows an illustration of the unloading of NLS or PR poly-dipeptide bound to Kapβ2 by RanGTP. **C**. Dose-dependent effect of PR20 expression on yeast growth. The percentage numbers indicate the galactose concentration used for induction of GFP-PR20 expression. **D**. Effect of PR20 expression on the motility of wild-type worms and *Cekapβ2* knockout. **E**. Turbidity of FUS droplets at different PR peptide concentrations in the absence and presence of Kapβ2. Droplet formation was triggered by cleavage of MBP-FUS by TEV protease. OD 395 nm is normalized to the measurement of MBP-FUS + buffer + TEV protease. Data are indicated as mean of three technical replicates ±SD. ** *p* < 0.01, * *p* < 0.05: statistical significance between the means of FUS turbidity in the absence and presence of Kapβ2 using a two-tailed *t*-test. n.d.: no significant difference. **F**. ODT images of FUS droplets in the presence of varying concentrations of PR20 in the absence and presence of Kapβ2. Scale bar, 5 µm. **G**. FRAP experiment of FUS droplet in the presence of Kapβ2 and excess PR20. Droplet formation was triggered by addition of 4% (w/v) PEG8000 to FUS solutions containing 9 µM MBP-FUS and 1 µM MBP-FUS-ATTO488 with or without 20 µM PR20 and 10 µM Kapβ2, and fluidity of FUS droplet assessed by FRAP. Fluorescence intensity before photobleaching was set to 100% and immediately after photobleaching was set to 0%. The change in relative fluorescence intensity was monitored for 120 s and calculated from raw data (Extended Data Fig. 5D). Data are indicated as mean of four‒six technical replicates + or −SD. Gray circle: FUS; red square: FUS, 2 eq PR20; purple triangle: FUS, 1 eq Kapβ2, 2 eq PR20. **H**. ODT and fluorescence imaging of FUS droplets with or without PR20 and Kapβ2. Droplet formation was triggered by addition of 4% (w/v) PEG8000 to FUS solutions containing 10 µM MBP-FUS and 10 µM Kapβ2-Alexa594 without (−) or with 10 µM (1 eq), 20 µM (2eq) PR20, respectively. Scale bar, 5 µm.

DPR toxicity to locomotion ability of *C. elegans* was evaluated based on the thrashing frequency. Expression of PR20 in motor neurons resulted in reduced thrashing frequency (Fig. 1D, Extended Data Fig. 4A), confirming the toxicity of DPRs, as previously shown in drosophila^26^ and mouse^27^. Interestingly, *C. elegans* lacking *imb-2*, a human Kapβ2 homolog in *C. elegans* (*Cekapβ2*), showed reduced thrashing frequency similar to that in WT expressing PR20. In addition, PR20 expression in *Cekapβ2*-deficient nematodes induced no further decrease in the thrashing frequency (Fig. 1D). This suggests that PR20 toxicity is expressed mainly through *Ce*Kapβ2. Indeed, direct binding between *Ce*Kapβ2 and PR20 was confirmed in vitro (Extended Data Fig. 4B and C). Additionally, motor neuron degeneration was seen in the nematodes expressing PR20, suggesting that PR20 impairs the locomotion ability of nematodes through motor neuron damages (Extended Data Fig. 4D). Notably, FUS was mislocalized in nematode neuron cells upon PR20 expression (Extended Data Fig. 4E and F) and *Cekapβ2* deletion^28^, implying that both PR20 expression and *Cekapβ2* deletion exert toxicity through dysregulation of RBPs.

## Excess PR poly-dipeptide turns Kapβ2 into a “phase disrupter”

A turbidity assay was performed to evaluate the effect of the excess DPRs on Kapβ2 activity as a phase modifier. In the assay, FUS droplet formation after removal of the solubility tag MBP by tobacco etch virus (TEV) protease cleavage was monitored at OD_395_ (Fig. 1E). While PR20 in the absence of Kapβ2 showed a negligible effect on turbidity, excess amount of PR20 significantly increased turbidity in the presence of Kapβ2, indicating a coexistence effect of PR20 and Kapβ2 on FUS droplet formation. SDS-PAGE analysis showed that the FUS droplets formed in the presence of Kapβ2 and 2 eq PR20 contained significant amount of Kapβ2 (Extended Data Fig. 5A). These data suggest that an excess PR20 not only inhibits Kapβ2 function as a phase modifier, but also triggers Kapβ2 accumulation in the FUS droplet.

To investigate the effect of Kapβ2 accumulation in the FUS droplet in the presence of excess PR poly-dipeptides in terms of the internal structure of the FUS droplet, 3D holographic RI imaging was performed. ODT observation showed that the internal density of FUS droplets, as indicated by RI, increased with the addition of excess PR20 only in the presence of Kapβ2 (Fig. 1F, Extended Data Fig. 5B, 6A, B, and Extended Data Table 1). Fluorescence recovery after photobleaching (FRAP) measurements of FUS droplets in the presence and absence of PR20 and Kapβ2 showed a distorted droplet shape and reduced fluorescence recovery in the presence of both Kapβ2 and 2 eq PR20, indicating reduced fluidity of FUS droplets (Fig. 1G and Extended Data Fig. 5C, D and 7). Fluorescence imaging of the FUS droplet in the presence of Alexa594-labeled Kapβ2 and PR20 showed PR20-dependent incorporation of Kapβ2 in the FUS droplet (Fig. 1H and Extended Data Fig. 6C). These results suggest that excess PR20 not only inhibits Kapβ2 activity as a phase modifier but turns it into phase disrupter, leading to the accumulation of PR20-bound Kapβ2 in the FUS droplet and altered properties of the droplet.

## Dose-dependent multiple binding of DPRs to Kapβ2

Dose-dependent interaction between Kapβ2 and DPRs at the molecular level was investigated using the “chemical tongue” technique, a biosensing technology that extracts multivariate pattern information reflecting sample characteristics by array of multiple fluorescent probes^29^. The chemical tongue can recognize a change in the exposed area of a protein due to complex formation^30^ (Extended Data Fig. 8A and B). In this study, complex formation between anionic Kapβ2 and cationic or neutral DPRs were monitored by using a set of cationic fluorescent polymers appended with aggregation-induced emission (AIE) luminogen, tetraphenylethylene (TPE)^31^ (Fig. 2A and Extended Data Fig. 8C–F). DPRs including PR20 at concentration range of 100 ∼ 6400 nM were mixed with 200 nM Kapβ2 in the presence of varying chemical probes (Fig. 2 and Extended Data Fig. 8G–I). PR20 and GR20, but not the other three DPRs, mixed with Kapβ2 induced changes in the fluorescence intensities, indicating the binding of PR20 and GR20 to Kapβ2 (Fig. 2A, Extended Data Fig. 8H and I), which is consistent with a previous study^19^. To extract more detailed information about the interaction, fluorescence response patterns were processed by using unsupervised principal component analysis (PCA)^32^ (Fig. 2B). On the PCA plot, the position of the clusters shifted significantly with the addition of PR20 and GR20 (Extended Data Fig. 8H). Interestingly, the change in the PCA score with increasing concentration of PR20 was biphasic (Fig. 2B). The PCA score (1) showed a rapid increase in the PR20 concentration range below ∼800 nM, whereas the score showed gradual increase in the higher concentration range (Fig. 2B middle panel). The PCA score (2) showed a rapid increase at lower concentration range, followed by a gradual decrease in the higher concentration range (Fig. 2B lower panel). These trends were also seen for GR20 (Extended Data Fig. 8J). Thus, chemical tongue analysis highlighted the existence of tight and weak interactions between Kapβ2 and DPRs.

**Figure 2.**
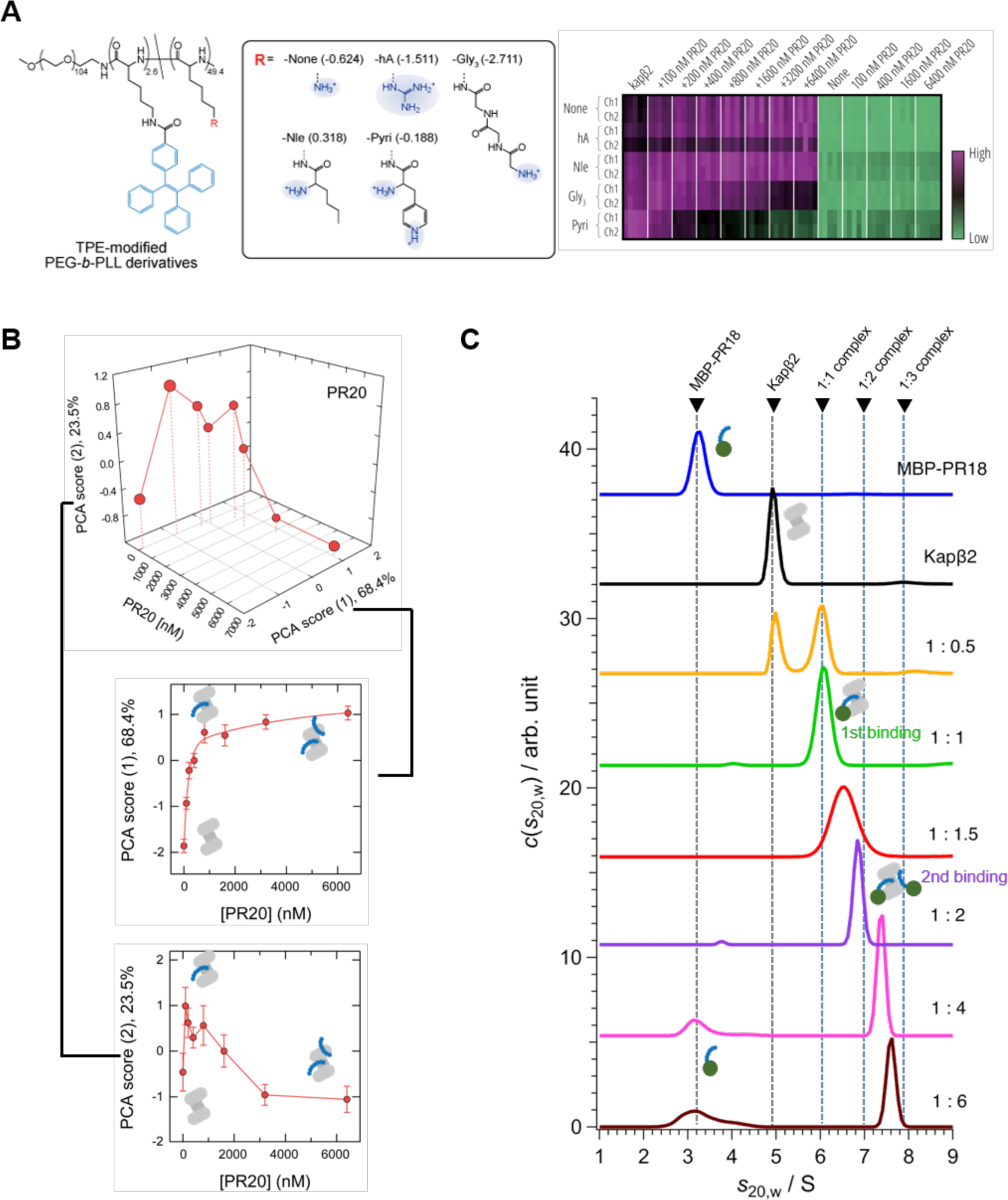
Chemical tongue assay showing the 2nd binding between Kapβ2 and PR poly-dipeptides. **A**. Cationic polymer probes used to extract interaction information between Kapβ2 and DPRs, and heat map of the fluorescence response patterns of Kapβ2 in the absence and presence of PR20. **B**. PCA scores for Kapβ2 in the absence and presence of PR20 plotted against PR concentration. **C**. AUC data for the solution of Kapβ2 and MBP-PR18 with various mixing ratio. Broken lines represent the sedimentation coefficient estimated from the molecular weight for each component.

Multiple interaction modes between DPRs and Kapβ2 were further corroborated by analytical ultracentrifugation (AUC) in which the size of the complex of MBP-PR18 and Kapβ2 was evaluated at varying molar ratios (Fig. 2C). The peak at 6.1 S observed below 1 equivalent mixing ratio (1:0.5 and 1:1) corresponds to Kapβ2:PR18 = 1:1 complex, suggesting tight binding at first site. As the mixing ratio of MBP-PR18 was increased beyond 1 equivalent (ranging from 1:1.5 to 1:6), the peak of the complex progressively shifted towards higher sedimentation coefficients (*s*_20,w_), suggesting the binding of multiple PR18 molecules to Kapβ2. Notably, the peak did not distinctly align with the *s*_20,w_ values expected for 1:2 or 1:3 complexes. This suggests that the rate of association and dissociation at the second binding site is considerably faster than the duration of the AUC measurement (four hours)^33^. Additionally, no peak corresponding to free MBP-PR18 appeared up to the addition of 2 equivalents of MBP-PR18 to Kapβ2, supporting that Kapβ2 can hold at least two molecules of PR18. Formation of Kapβ2:PR18 = 1:2 complex was also detected in the analytical gel filtration (Extended Data Fig. 9). In support of PR poly-dipeptides binding to multiple sites on Kapβ2, hydrogel binding experiments showed that a sub-stoichiometric amount of Kapβ2 inhibited the binding between PR20 and FUS LC (Extended Data Fig. 10).

Collectively, the chemical tongue analysis, AUC, analytical gel filtration, and hydrogel binding assay indicated that the interaction between Kapβ2 and PR poly-dipeptides consists of tight and weak binding modes. The weak binding mode may be a key to understanding the mechanism of dose-dependent toxicity of DPRs.

## The conserved acidic loop of Kapβ2 is responsible for weak binding with DPRs

Previously, we have shown that DPRs tightly bind Kapβ2 at its NLS binding site, which consists of a negatively charged cavity^19^. This was further supported by the chemical tongue analysis in this study. In addition to the NLS binding site, Kapβ2 has a negatively charged 51-residue loop containing 24 acidic amino acids (Fig. 3A and Extended Data Fig. 11), which we suspected could be the second binding site of DPRs on Kapβ2. The abundance of acidic residues in the loop is highly conserved from yeast to human (Fig. 3B). To test whether PR poly-dipeptides directly interacts with the conserved acidic loop of Kapβ2, we prepared a ^15^N-labeled Kapβ2 loop whose N-terminus is fused to protein G B1 domain (GB1) T19C mutant and the C-terminus fused to additional Cys residues to form disulfide-bond with GB1 T19C (Fig. 3C). ^1^H-^15^N hetero-nuclear single quantum coherence (HSQC) spectra, in the absence and presence of PR20, showed that PR20 induced significant chemical shift changes and peak broadening to the several resonances from the loop (Fig. 3C, D, and Extended Data Fig. 12), showing the direct interaction between PR20 and the Kapβ2 loop.

**Figure 3.**
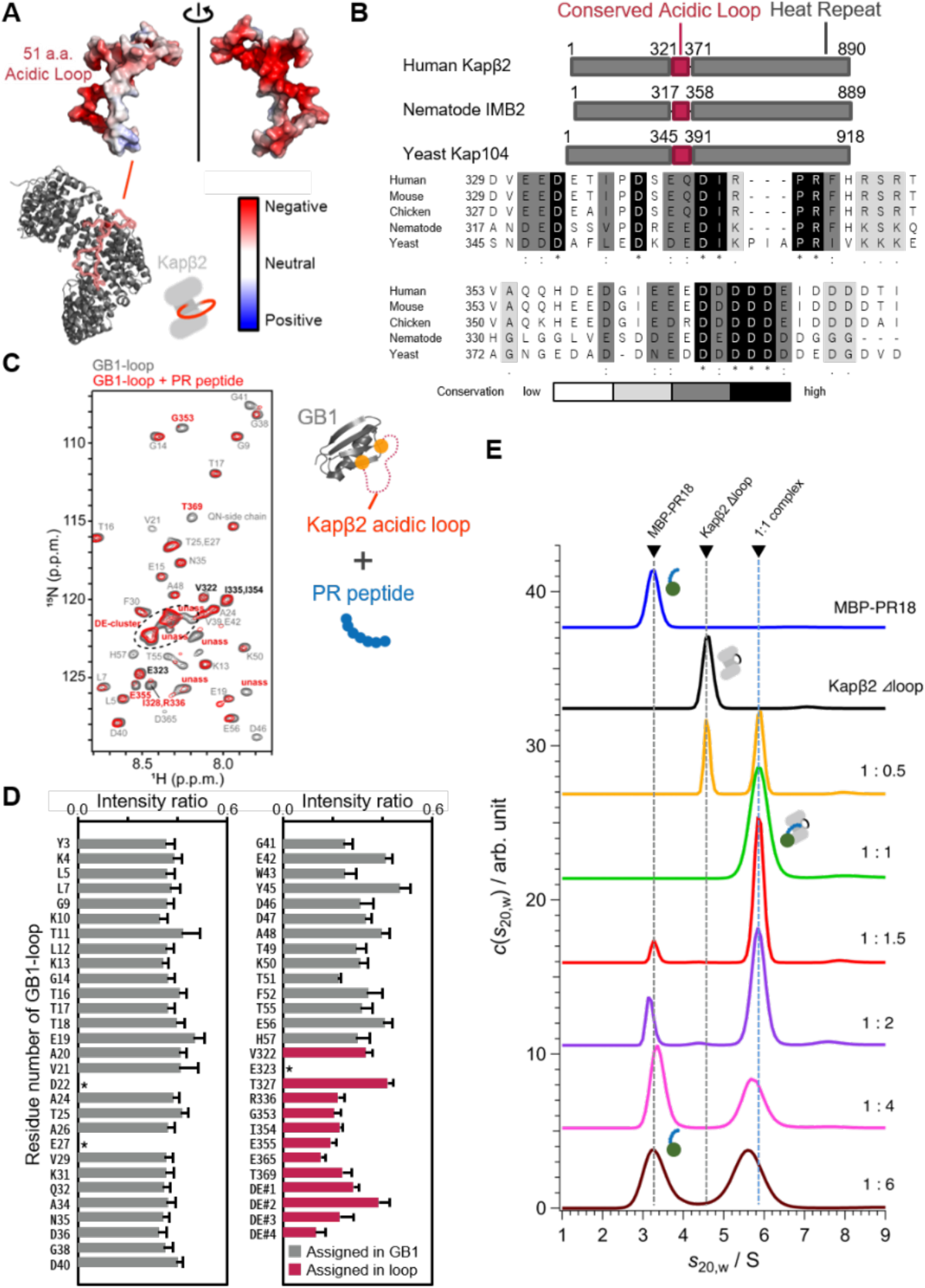
The loop region of Kapβ2 constitutes the 2nd binding site of PR poly-dipeptides. **A**. Predicted structural model with surface electrostatic potential of the Kapβ2 loop region and crystal structure of Kapβ2. The crystal structure (PDB code: 2z5j) is shown in gray cartoon. The acidic loop consisting of 51 residues is shown in red lines. **B**. The schematic representation of Kapβ2 and its homologs and multiple alignment of their conserved acidic loops. The full length of Kapβ2 homologs is indicated by a gray box and the acidic loops within it by a red box. Alignment of the amino acid sequence of conserved acidic loops was made from the genes of Kapβ2 homologs in each species (human: NP_002261.3, mouse: NP_848831.2, chicken: XP_424806.3, nematode: NP_496987.1, yeast: NP_594385.1). Highly conserved residues are highlighted in gray and black boxes. **C**. ^1^H-^15^N HSQC spectra of Kapβ2 loop fused to GB1 “GB1-loop” in the absence (gray) and presence (red) of PR20. The regions indicated by a box in the full spectra (Extended Data Fig. 12) are expanded with resonance assignments. The resonances showing significant perturbations are indicated by broken line. The “GB1-loop” consists of GB1 T19C mutant and the conserved acidic loop sequence of Kapβ2, and C-terminal of the loop sequence are linked by using disulfide bond to structurally mimic the acidic loop of Kapβ2. **D**. Plots of peak intensity change of GB1-loop by addition of PR20. The intensity ratio of resonances assigned in GB1 region (gray) and assigned in loop region (red) are indicated as mean +SD estimated from noise level. The resonances disappeared by the addition of PR20 are indicated asterisks. **E**. AUC data for the solution of Kapβ2 Δloop and MBP-PR18 with various mixing ratio. Broken lines represent the sedimentation coefficient estimated from the molecular weight for each component.

Interaction between PR20 and the loop was corroborated by AUC analysis on Kapβ2 Δloop and MBP-PR18. In this analysis, the 1:1 complex was observed even in the presence of an excess amount of MBP-PR18 along with a peak corresponding to the unbound MBP-PR18 (Fig. 3E, 1:1.5 ∼ 1:6), indicating that Kapβ2 Δloop cannot hold more than 1 equivalent of PR18. This suggests that the conserved acidic loop has a critical role in the binding between Kapβ2 and PR poly-dipeptides. Most of the resonances in the NMR spectrum of Kapβ2 Δloop matched with those from Kapβ2 WT, with few exceptions corresponding to the resonances derived from the loop (Extended Data Fig. 13), confirming that the overall structure of Kapβ2 is retained in the Δloop mutant. In addition, the affinity of Kapβ2 Δloop to PR20 labeled with a fluorophore dansyl group (Dnc-PR20) was estimated to be *K*_D_ = 3.1×10^2^ nM (Extended Data Fig. 14), suggesting that the 1st binding site for PR, the NLS-binding site^19^, is also preserved in the Kapβ2 Δloop. Collectively, our data showed that PR poly-dipeptides interacts not only with the NLS binding site but also with the conserved acidic loop of Kapβ2 as a 2nd binding site.

## The conserved acidic loop of Kapβ2 plays an essential role in phase modification

Our experiments demonstrate that Kapβ2 dysfunction and retention inside the droplet is caused by the 2nd binding of PR. This implies that the conserved acidic loop provides an essential function of Kapβ2 as a phase modifier. To investigate the function of the loop, we performed RI imaging for Kapβ2 Δloop-Alexa594 (Fig. 4A; Extended Data Fig. 15A). The results showed that Kapβ2 Δloop was incorporated into the FUS droplet even without PR poly-dipeptides, and the density of the droplet was increased (Fig. 4A and B). FRAP measurements showed that the addition of Kapβ2 Δloop to the FUS droplet decreased the fluidity inside the droplets (Fig. 4C and Extended Data Fig. 15B and C). These observations clearly show that the conserved acidic loop is essential for maintaining the chaperone ability of Kapβ2.

**Figure 4.**
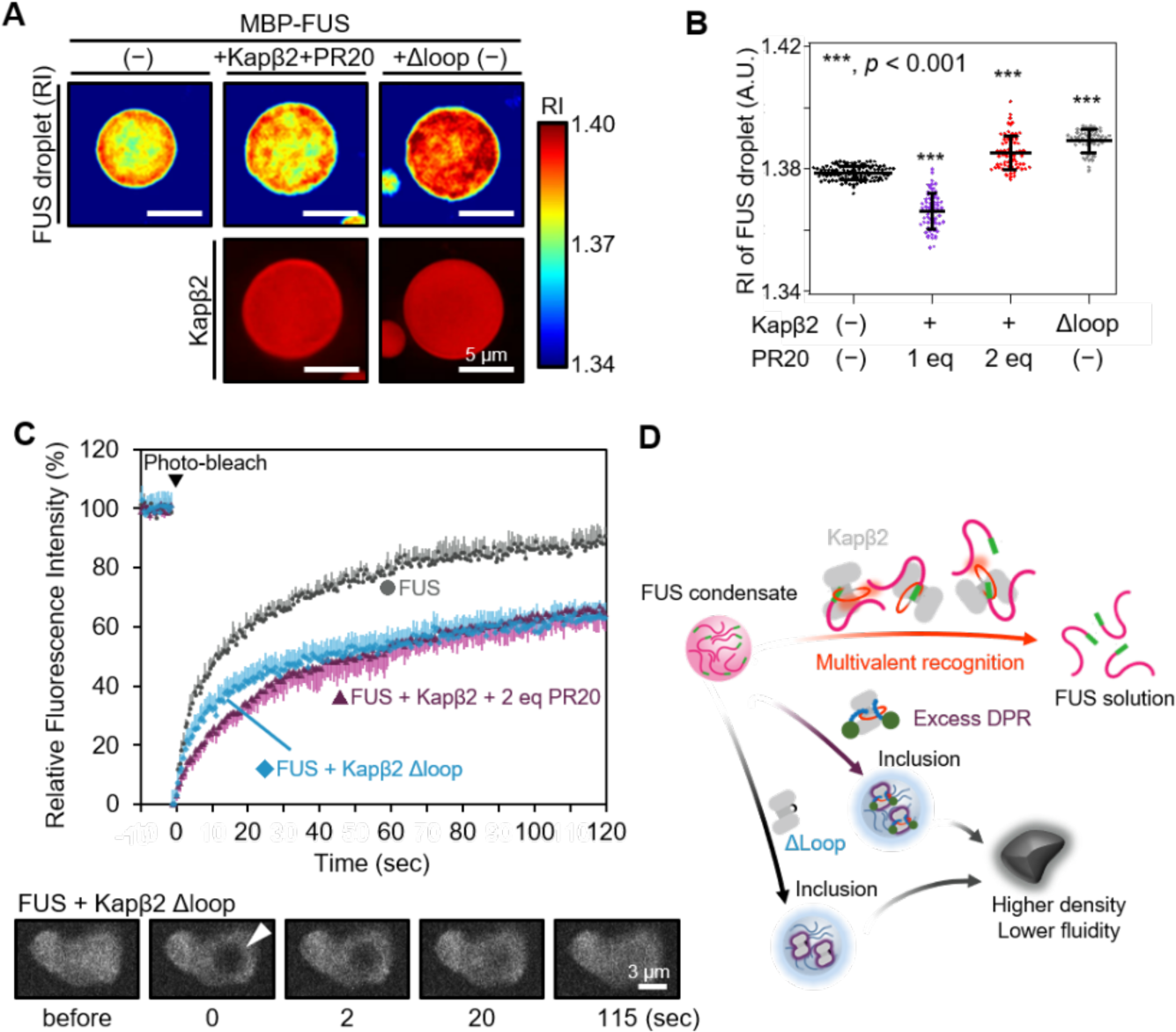
The excess PR poly-dipeptides inhibits Kapβ2 as a phase modifier and promotes aggregation of Kapβ2 and FUS in the FUS droplet. **A**. ODT and fluorescence images of FUS droplets in the absence and presence of Kapβ2 and 2 eq PR20 or Kapβ2 Δloop. Droplet formation was triggered by addition of 4% (w/v) PEG8000 to FUS solutions containing 10 µM MBP-FUS without (−), with 10 µM Kapβ2-Alexa594 and 20 µM (2eq) PR, or with 10 µM Kapβ2 Δloop-Alexa594. Scale bar, 5 µm. **B**. Quantitative analysis of RI data shown in **A**. Data are mean±SD. MBP-FUS, *n* = 113 droplets from five independent experiments; MBP-FUS + Kapβ2 + 1 eq PR20, *n* = 66 droplets from five independent experiments; MBP-FUS + Kapβ2 + 2 eq PR20, *n* = 74 droplets from six independent experiments; MBP-FUS + Kapβ2 βloop, *n* = 59 droplets from three independent experiments. **C**. FRAP experiment of FUS droplet in the absence and presence of Kapβ2 and 2 eq PR20 or Kapβ2 Δloop. Experimental condition is the same as that in Figure 1G. The change in relative fluorescence intensity was monitored for 120 s and calculated from raw data (Extended Data Fig. S4-1A and B). Data are indicated as mean of four‒six technical replicates + or −SD. Gray circle: FUS; blue diamond: FUS, 1 eq Kapβ2 Δloop, purple triangle: FUS, 1 eq Kapβ2, 2 eq PR20. **D**. Schematic representation of molecular mechanism behind the irreversible breakdown of the Kapβ2 phase-separation regulation. Free Kapβ2 is involved in LLPS regulation of FUS (red arrow). When excess DPRs are present, DPRs bind to multiple sites, including the NLS binding site and conserved acidic loop of Kapβ2. Multiple DPR-bound Kapβ2 loses its function to regulate LLPS via the inhibitory interaction between the acidic loop and DPRs and is then incorporated into FUS droplets. Kapβ2 Δloop mutation has a similar effect and disrupts the phase modifier-function of Kapβ2. In the end, FUS droplets with insoluble and malfunctional Kapβ2 irreversibly become denser and less fluid.

The above functional analyses of Kapβ2 with excess DPRs suggest that the 2nd binding of DPRs following the 1st binding to the NLS binding site is the initiation of Kapβ2 dysfunction and that the incorporation of the Kapβ2 and multiple-DPR complex into the droplet leads to the irreversible breakdown of the Kapβ2 phase-separation regulation (Fig. 4D).

## Discussion

DPRs produced from *C9orf72* repeat expansion is highly neurotoxic, but the mechanism of toxicity is not fully understood. A key pathway of DPR toxicity is through the nuclear transport receptor Kapβ2. We have previously shown that the tight binding of DPRs to the NLS-binding site of Kapβ2 inhibits its interaction with RBPs and subsequently inhibits its activity as a phase modifier^19^. Our previous study, as well as other recent studies^34–36^, suggest that DPRs can bind to regions of Kapβ2 other than the NLS-binding site, but the actual interaction sites remained to be elucidated. This study identified the conserved acidic loop of Kapβ2 as a second binding site for DPRs (Fig. 2 and 3). This is consistent with a recent simulation study that shows the binding of DPRs to multiple regions of Kapβ2 including the NLS-binding site, loops, and outer surface areas^34^. Intriguingly, the PR poly-dipeptides binding to the Kapβ2 loop not only inhibits the phase modifier activity of Kapβ2, but also alters it to a “phase disrupter”, leading to the accumulation of Kapβ2 in the FUS droplet, which, in turn, makes the droplet more condensed with decreased internal mobility, possibly leading to insoluble aggregates (Fig. 4). Assays using Kapβ2 Δloop showed that the loop is necessary for Kapβ2 to work as a phase modifier, highlighting the importance of multivalent recognition of RBPs by the Kapβ2 loop to dissolve RBD condensates. The acidic loop of Kapβ2 has also been shown to be responsible for the recognition of RGG domain of CIRBP^39^. Given that the RBPs also possess RGG domain, the recognition of RGG domain by the Kapβ2 loop should be critical as a phase modifier. Oligomerization of PR-bound Kapβ2, as shown by previous studies^35,38^ may also contribute to the accumulation of Kapβ2 in FUS droplet. Incorporation of Kapβ2 into the RBP condensates can reduce the amount of active Kapβ2, resulting in the disruption of the nuclear transport pathway and RBP accumulation in cytosol.

Moreover, our data showed that DPRs ride on Kapβ2 to be transported and accumulated in the nucleus (Fig. 1), which explains the mechanism of the PR poly-dipeptide accumulation observed in the *C. elegans* model and yeast cells (Fig. 1) as well as in a monkey model^37^. Although we showed multiple binding of DPRs to Kapβ2 using dipeptides with 18 or 20 repeats, we expect longer DPRs as a single chain to be able to simultaneously bind multiple regions of Kapβ, including the NLS-blinding site and the acidic loop. This explains the higher toxicity of longer repeat expansion at *C9orf72*^1^.

This study has revealed the mechanism by which DPRs accumulate in the nucleus and exert toxic effects. Structural and biochemical approaches also revealed the functional importance of the conserved acidic loop as the phase modifier, and binding of multiple DPRs molecules to the loop reverses its function. It also suggests that RBPs and Kapβ2 are key players in the background of familial ALS caused by DPR toxicity and further clarification of the relationship between the mechanism of DPR toxicity and RBP pathology is expected to elucidate the mechanism of familial ALS onset and progression.

## Acknowledgements

This work was supported by funding from JSPS KAKENHI (JP21J21141/JP22KJ0024 to H.Ka., JP20K15969, JP22H04847, and JP22K15278 to M.M., JP19H04945, JP20H03199, JP20KK0156, JP21H05093, JP21H05094, JP22H02560, JP22K18361, JP23H05470, and JP23H01995 to T.S., JP23H01995 to S.T., JP20K06493 to T.Man., JP23H01336 and JP23H04396 to T.Mab.), MEXT Grant-in-Aid for Transformative Research Areas (B) (JP21H05094 and JP21H05093 to T.S., JP21H05096 to T.Mab., JP21H05095 to M.O.), AMED (JP24ek0109642 to T.S., Y.S., T.N., JP21ek0109558 to H.N., T.Man., T.Y., JP24wm0425004 to E.M., T.S., S.T., T.N.), AMED-BINDS (JP22ama121001 to M.S.), JST FOREST Program (JPMJFR212H to T.Mab., JPMJFR201F to M.O., and JPMJFR204W to T.S.), HIRAKU-Global Program, which is funded by MEXT’s “Strategic Professional Development Program for Young Researchers” to M.M, and the Tohoku Initiative for Fostering Global Researchers for Interdisciplinary Sciences (TI-FRIS) of MEXT’s Strategic Professional Development Program for Young Researchers to T.Mab. This work was also partially supported by Astellas Foundation for Research on Metabolic Disorders and The Uehara Memorial Foundation to M.M., Takeda Science Foundation Grant, The Sumitomo Foundation, Astellas Foundation for Research on Metabolic Disorders, Senri Life Science Foundation, The Nakajima Foundation, The Asahi Glass Foundation, Akiyama Life Science Foundation Grants-in-Aid, Northern Advancement Center for Science and Technology Grants-in-Aid, The Nakabayashi Trust For ALS Research, The Kato Memorial Trust for Nambyo Research, Mochida Memorial Foundation for Medical and Pharmaceutical Research, and The Naito Foundation to T.S. This work was performed in part using the NMR spectrometers with the ultra-high magnetic fields under the Collaborative Research Program of Institute for Protein Research, Osaka University, NMRCR-2023-05, and was supported by the program of the Inter-University Research Network for High Depth Omics, and Joint Usage and Joint Research Programs, the Institute of Advanced Medical Sciences (IAMS), Tokushima University. This research in part used experimental resources provided by Fujii Memorial Institute of Medical Sciences, IAMS, Tokushima University. AUC experiment was performed at Institute for Integrated Radiation and Nuclear Science, Kyoto University (KURNS) under Proposal Number R4129. We also thank the Bio-support Center at Tokyo Tech for DNA sequencing, and the Cell Biology Center Research Core Facility at Tokyo Tech for providing the IXplore SpinSR microscope system. Some *C. elegans* strains were provided by NBRP, which is funded by the Japanese government, and the CGC, which is funded by NIH Office of Research Infrastructure Programs (P40 OD010440). We thank Dr. Steven L. McKnight (University of Texas Southwestern) for providing mCherry fusion LC-domain of FUS and GFP fusion 20 repeats of PR constructs.

## Author contributions

E.M., T.Y., and T.S. conceived the research project and the overall design of the experimental strategy. H.Ka., H.Ku., and T.S. performed NMR experiments. H.Ka., M.M., K.K., Y.H., T.Mu., and T.S. performed FRAP experiments and fluorescence assays. T.T., T.U., H.M., and T.Y. performed turbidity assay and analytical SEC. M.W., S.K., and M.O. performed ODT experiments. K.M. and M.S. performed AUC experiments. S.T. performed chemical tongue experiments. Y.S. performed *C. elegance* in vivo experiments. T.N. performed yeast in vivo experiments. T.Man., N.I., and E.M. performed the cell-based experiments. H.N., K.S., and E.M. performed hydrogel experiments. T.Mab. and Y.A. performed structural modeling. K.I. and T.Mu. provided critical intellectual input and data interpretation. H.K., M.M., and T.S. prepared the original draft with input from all authors. All authors discussed the results and commented on the manuscript.

## Data availability

The data supporting the findings of this study are available from the corresponding authors upon reasonable request. Source data are provided with this paper.

## Methods

### Antibodies and reagents

Antibodies were procured from the following sources: anti-HA (Cat. No. 51064-2-AP; proteintech), anti-KDEL (Cat. No. M181-3; Medical & Biological Laboratories (MBL) Co., Ltd., Nagoya, Japan), anti-c-Myc (Cat. No. sc-40; Santa Cruz Biotechnology, Dallas, TX), anti-c-Myc (Cat. No. 562; MBL), anti-GAPDH (Cat. No. G9295; Sigma–Aldrich, St. Louis, MO), anti-PDI (Cat. No. ADI-SPA-891; Enzo Life Sciences, Farmingdale, USA), and anti-HA agarose (Cat. No. #11815016001, Roche, Switzerland). PR20 modified with a dansyl group at the N-terminus (Dnc-PR20; ≥95% purity) was purchased from Biologica Co. (Nagoya, Japan).

### Expression and purification of protein samples

Kapβ2 loop (321‒371) expression construct was cloned into pET21b vector (Cat. no. 69741-3CN; Novagen, Madison, Wisconsin, USA) and fused to GB1-His^6^ tags. The crystal structure of Kapβ2 (PDB code 2Z5J) shows that the distance between Asp321 and Glu371 is ∼12. The GB1-loop distance between Thr44 and the terminal of the loop was ∼12 Å. We introduced a T44C mutation and 108C mutation into the GB1-loop by site-directed mutagenesis using a PrimeSTAR mutagenesis basal kit (Takara Bio).

The ^15^N-labelled GB1-loop protein was overexpressed in *E. coli* BL21 (DE3) cells, which were grown in M9 minimal medium with ^15^NH_4_Cl (CIL) as the sole nitrogen source and induced with 0.5 mm IPTG at OD_600_ ∼ 0.6, followed by ∼16 h of incubation at 25 °C. Cells were collected and re-suspended in the lysis buffer containing 50 mM Tris-HCl pH 8.0, 500 mM NaCl. Cells were disrupted by a sonicator and centrifuged at 15,000 rpm for 30 min. Proteins were purified by Ni-NTA affinity chromatography (QIAGEN) and gel filtration chromatography (HiLoad 26/60 Superdex 75 pg, GE Healthcare). Finally, the construct was diluted to a concentration of 10–20 μM and incubated with 1 mM 5,5′-dithiobis(2-nitrobenzoic acid) (DTNB) for 2 h at room temperature to form an intramolecular disulfide bond. After incubation, DTNB was removed by dialysis.

Unlabeled recombinant proteins for biochemical and biophysical analyses were expressed individually in BL21(DE3) cells^19^. Kapβ2 was expressed from a GST-fusion with HRV3C protease site in between the GST and Kapβ2. Kapβ2 was purified using Glutathione sepharose beads (Cytiva); the GST was cleaved by HRV3C protease, followed by anion exchange chromatography (HiTrap Q HP, Cytiva), gel filtration chromatography (Superdex200 16/60, Cytiva) using a buffer containing 20 mM Hepes pH 7.4, 150 mM NaCl, 2 mM DTT, 2 mM Mg(OAc)_2_, and 10% Glycerol. MBP-fused proteins, MBP-PR18, MBP-TEV-FUS, and MBP-hnRNPA2 (NLS, residues 298‒336) were purified using affinity chromatography (Amylose resin, NEB), and MBP-PR18 was further purified using cation exchange chromatography (HiTrap SP HP, Cytiva) and gel filtration chromatography (Superdex200 16/60, Cytiva). GST-fused GTP-bound Ran (Q69L), RanGTP was purified using GSH sepharose beads (Cytiva); the GST was cleaved by HRV3C protease, followed by anion exchange chromatography (HiTrap Q HP, Cytiva) and gel filtration chromatography (Superdex200 16/60, Cytiva).

### Evaluation of DPR localization in cultured mammalian cells

The pcDNA5/FRT/TO-HA-SBP-GFP-PR20-HA plasmid (GFP-PR-HA) was constructed as described previously^19^. The pcDNA5/FRT/TO-HA-SBP-GFP-HA plasmid (GFP-HA) was generated by deleting the PR20 sequence using PCR-based site-directed mutagenesis of the GFP-PR-HA plasmid. The pcDNA3-5’-Venus (Venus) plasmid was constructed as described previously^40^. The pcDNA3-5’-Venus-M9M plasmid was generated by inserting the M9M fragment into the Venus plasmid between the *Bam*HI and *Not*I sites. HeLa cells were cultured in DMEM/10% FBS at 37 °C with 5% CO_2_. For microscopic observation, cells were seeded onto round, 12-mm-diameter coverslips of 24-well plates and transfected with plasmid using TransIT LT1, according to the manufacturer’s instructions. The cells were fixed 24 h after transfection with 4% paraformaldehyde prepared in PBS at room temperature (RT) for 10 min. The fixed cells were permeabilized with 0.5% Triton X-100 prepared in PBS for 15 min, rinsed, and blocked with 1% BSA prepared in PBS containing 0.1% Tween-20 (PBST) for 1 h. The slides were incubated at 4 °C overnight with primary antibodies (diluted in PBST containing 1% BSA) against specific proteins. Unbound antibodies were removed with three 10 min washes with PBST. The slides were then incubated with Alexa-conjugated secondary antibodies for 1 h at RT, washed, and mounted with Fluoro-KEEPER Antifade Reagent (Nacalai Tesque). Immunostained cells were examined using a confocal laser scanning microscope (FV1000D and FV10i; Olympus).

### In vitro pull-down assays

In vitro pull-down binding assays showing the interaction between MBP-PR18 and Kapβ2 were performed using GST-Kapβ2 immobilized on GSH sepharose beads (GE Healthcare). Briefly, 10 μM GST-Kapβ2 were immobilized on 30 μL of beads. The GST-proteins beads were incubated with 20 μM proteins, either MBP-PR18 or MBP fused with the NLS region of hnRNPA2 (NLS, residues 298-336) for 20–30 min and washed three times with a buffer containing 20 mM Hepes pH 7.4, 150 mM NaCl, 10% glycerol, 2 mM Mg(OAc)_2_, and 2 mM DTT. For the competitive binding assay, RanGTP was added to the immobilized beads and unbound proteins were washed three times with the buffer. Thereafter, bound and unbound proteins were separated by SDS-PAGE and stained with Coomassie Brilliant Blue.

### Yeast cell assays

*Saccharomyces cerevisiae* (Baker’s yeast) strain (BY4741: *MATa his3 Δ1 leu2 Δ0 met15 Δ0 ura3 Δ0*) harboring the YEplac plasmids (vacant vector or containing GFP, GFP-PR8, or GFP-PR20 gene controlled by *GAL1* promoter) were grown in SRaf-Ura liquid medium, a synthetic complete medium containing 2% raffinose as a carbon source and lacking uracil, to the late logarithmic growth phase.

For the growth curve assay, cells were diluted to 0.002 OD_600_ and cultured at 30 °C with shaking (70 rpm) under four carbon source conditions: 2% raffinose (0% Gal), 2% raffinose and 0.02% galactose (0.02% Gal), 1.8% raffinose and 0.2% galactose (0.2% Gal), and 2% galactose (2% Gal). During the culture, OD_600_ was measured continuously every 30 min using a biophotorecorder (TVS062CA, Advantec Toyo Co Ltd., Tokyo, Japan).

For the spot assay, the cell culture was serially diluted (10^7^, 10^6^, 10^5^, 10^4^, and 10^3^ cells/mL), and 4 μL of the cell suspensions was spotted on an SRaf-Ura agar plate containing 2% glucose or 2% galactose. The spotted plates were incubated at 30 °C for two to three days.

For the microscopic observation, the late log cultures were diluted 500–1000 times and cultured again in SRaf-Ura liquid medium to around 0.6 OD_600_. Galactose was then added to a final concentration of 0.2%, and the cells were cultured for 3 h for induction. After induction, Hoechst 33342 (Dojindo, Kumamoto, Japan) was added at a final 1 µg/mL concentration and cultured further for 10 min. For the visualization of GFP and Hoechst 33342 fluorescence in vivo, the IXplore SpinSR microscope system (Evident, Tokyo, Japan) was used in the confocal mode. The objective lens used was UPlanApo 100×/1.50 NA oil immersion (Evident, Tokyo, Japan), and the ORCA Flash 4.0 CMOS camera (Hamamatsu Photonics, Shizuoka, Japan) was used as a detector. The obtained pictures were processed using ImageJ software (https://imagej.net/ij/).

### C. elegans assays

All the strains used in this study were wild-type strain N2 and *imb-2(tm6405)/mnC1[nIS190]*. All strains were cultured on NGM plates with *E. coli* strain OP50 at 20 °C. Transgenic strains were generated by microinjecting *Punc-47::PR20::SL2::wrmScarlet* construct or *Punc-47::egfp::fus*. The fluorescence image of adult-stage hermaphrodites harboring *Punc-47::PR20::SL2::wrmScarlet, Punc-47::egfp::fus*, or *Punc-25::gfp* was obtained with an Axioplan 2 microscope (Zeiss), a FV1000 laser-scanning confocal microscope (Olympus), or a THUNDER microscope (Leica). Thrashing behavior was measured using adult day 1 worms suspended in S-basal buffer and analyzed by WormLab software (MBF Bioscience). Since *imb-2(tm6405)* homozygous mutants grow to adulthood and show maternal lethal phenotype, they were produced from *imb-2(tm6405)/mnC1[nIS190]* heterozygous mutants. Statistical significance was determined by Student’s *t*-test (p < 0.01).

### FUS turbidity assay

Prior to adding TEV protease, 8 μM MBP:FUS, ± 8 μM Kapβ2 and ± PR20:HA were mixed in buffer containing 20 mM Hepes pH 7.4, 150 mM NaCl, 10% glycerol, 2 mM Mg(OAc)_2_, 20 μM Zn(OAc)_2_, and 2 mM DTT to reaction volumes of 100 μL. TEV protease was added to the premixture to a final concentration of 40 μg/mL and incubated at 30 °C for 60 min to digest all MBP-fusion proteins. The solution was left to cool down to 20 °C before measurement of OD at 395 nm using a plate reader (SH1000, CORONA). The mixtures were centrifuged (16,441 *g*, 10 min, 4 °C) and then proteins in the supernatant and precipitant were separated by SDS-PAGE and stained with Coomassie Brilliant Blue. Statistical significance was determined by a two-tailed Welch’s *t*-test (p < 0.05) using Microsoft Excel software (Microsoft, Roselle, IL, USA).

### Hydrogel binding assay

Expression plasmids for recombinant proteins (mCherry fusion LC-domain of FUS [residue 2–214, mCh:FUS-LC] and GFP fusion 20 repeats of PR [GFP:PR20]) were obtained from the Steven L. McKnight Laboratory. mCh:FUS-LC and GFP:PR20 were expressed in *E. coli* BL21(DE3) cells and purified by Ni-NTA Agarose (FUJIFILM Wako Pure Chemical Corporation), as described in a previous study^41^. Hydrogel droplets of mCh:FUS-LC were prepared as reported in a previous study^41^. For hydrogel binding assays, purified GFP:PR20 was diluted to 0.5 µM in the buffer (20 mM Tris-HCl pH 7.5, 150 mM NaCl, 20 mM β-mercaptoethanol, 0.1 mM phenylmethylsulfonyl fluoride, and 0.5 mM EDTA) and pipetted onto a hydrogel dish. After overnight incubation, horizontal sections of the droplets were scanned on a confocal microscope (FLUOVIEW FV3000, OLYMPUS).

### RI measurement inside the FUS droplet using ODT

The LLPS-droplet formation was initiated by mixing 10 μM of MBP-FUS with 4% polyethylene glycol 8000 in a buffer containing 20 mM HEPES-NaOH, pH 7.4, 150 mM NaCl, 10% glycerol, and 2 mM DTT. After 20 min, 10 μM of Alexa594-Kapβ2 and/or 10 or 20 μM of PR20 was added to the mixture and incubated for 17 min. The droplets were observed using a holotomography microscope with laser-induced fluorescence system^42,43^ (Tomocube, Inc.). The RI and fluorescence image show the XY cross section of the droplet center. The average RI inside the droplet and their radius were calculated by enclosing the RI image in a circle using TomoStudio version 3.2.8 software (Tomocube, Inc.) and plotted using the Igor Pro version 6.36 software (WaveMetrics). The statistical significance of differences was examined by one-way analysis of variance with Tukey honestly significant difference (HSD) post hoc testing. All statistical tests were performed using KaleidaGraph version 4.5.1 software (Synergy Software) at a significance level of α = 0.05.

### Fluorescence recovery after photobleaching

Droplet formation was triggered by addition of 4% (w/v) PEG8000 to FUS solutions containing 9 µM MBP-FUS, 1 µM MBP-FUS-ATTO488, 20 mM HEPES-NaOH pH 7.4 at room temperature, 150 mM NaCl, 2 mM DTT, and 10% (w/v) glycerol in the absence or presence of 20 µM PR20, and the FUS solution was well-mixed and immediately dropped onto a glass bottom 10-compartment cell culture slide (CELLview, Cat. #543079, Greiner Bio-One, Kremsmünstern, Austria). FUS droplets were observed with the 473 nm laser line of a confocal microscope (FV1200, Olympus, Tokyo, Japan) that was equipped with a UPLSAPO40X2 objective lens (NA 0.95). A specific spot in the FUS droplet was bleached with 100% transmission at 20th and 21st frames, and time-lapse images before and after photobleaching were collected (0.5 frames per second, 260 frames). Fluorescence intensity of the region of interest was then calculated using FV10-ASW software (Olympus). The images were processed using ImageJ/Fiji^44^ and GIMP (GNU Image Manipulation Program) software (http://gimp.org). Fluorescence intensity before photobleaching was set to 100% and immediately after photobleaching was set to 0%, and half time of fluorescence recovery and apparent diffusion coefficients were calculated from the normalized fluorescence intensity using Microsoft Excel (Microsoft), based on previous studies^45–47^.

### Interaction analysis using chemical tongue

#### Fluorescence response of the polymer probes

Polyethylene glycol-*block*-poly-L-lysine (PEG-*b*-PLL) modified with tetraphenylethylene (TPE) and their derivatives were synthesized according to a previously reported method^31^. Fluorescence measurements were performed using a Cytation5 Imaging Reader (BioTek Instruments, Inc.). Solutions (60 μL) containing 100 nM TPE-functionalized PEG-*b*-PLLs, analytes (0–500 nM for kapβ2; 0–2000 nM for DPRs), 150 mM NaCl, and 2 mM DTT in 20 mM Tris-HCl buffer (pH = 7.5) were prepared in each well of a 384-well NBS^TM^ black microplate (Corning Inc.) using an Andrew+ liquid handling robot (Andrew Alliance SA, Geneva, Switzerland). After incubation (35 °C, 10 min), the fluorescence spectrum (λ_ex_/λ_em_ = 330 nm/372–700 nm) or the fluorescence intensity (λ_ex_/λ_em_ = 330 nm/480 nm) were recorded at 35 °C. To assess the interaction between Kapβ2 and DPRs, solutions (60 μL) containing 100 nM TPE-functionalized PEG-*b*-PLLs, 200 nM kapβ2, DPRs (0–1600 nM), 150 mM NaCl, and 2 mM DTT in 20 mM Tris-HCl buffer (pH = 7.5) were prepared and analyzed as well. The displayed values represent mean values ± standard error of mean from three independent experiments.

#### Chemical tongue sensing

Fluorescence measurements were performed using a Cytation5 Imaging Reader. For the evaluation of concentration-dependence of Arg-containing DPRs, aliquots (20 μL) of solutions containing TPE-functionalized PEG-*b*-PLLs (150 nM), 225 mM NaCl, and 3 mM DTT in 30 mM Tris-HCl buffer (pH = 7.5) were deposited in the wells of a low volume 384-well plate (Corning Inc.) using an Andrew+ liquid handling robot. After incubation (35 °C, 10 min), the fluorescence intensity was recorded using two different channels (Ch1: λ_ex_/λ_em_ = 330 nm/480 nm; Ch2: λ_ex_/λ_em_ = 360 nm/530 nm). Subsequently, aliquots (5 μL) of kapβ2 and aliquots (5 μL) of DPRs were added to each well, and the fluorescence intensity was recorded after incubation (35 °C, 10 min). Final concentrations were 100 nM TPE-functionalized PEG-*b*-PLLs, 0 or 200 nM kapβ2, 0–6400 nM DPRs, 150 mM NaCl, and 2 mM DTT in 20 mM Tris-HCl (pH = 7.5). For the comparison of DPRs, measurements were similarly conducted with final concentrations of 100 nM TPE-functionalized PEG-*b*-PLLs, 0 or 200 nM kapβ2, 0–600 nM DPRs, 150 mM NaCl, and 2 mM DTT in 20 mM Tris-HCl (pH = 7.5). These processes were performed at least five times for distinct samples. This dataset was processed using PCA in SYSTAT 13 (Systat Inc.). The displayed ellipsoids represent the confidence intervals ± standard deviations for each analyte.

### Analytical ultracentrifugation (AUC)

Samples were prepared in a buffer containing 20 mM HEPES-NaOH, pH 7.4, 150 mM NaCl, 10% glycerol, 2 mM DTT, and 2 mM magnesium acetate. The concentrations of Kapβ2 and Kapβ2 Δloop were fixed at 28 μM for the titration measurement with various mixing ratio.

AUC measurement was performed using ProteomeLab XL-I (Beckman Coulter). Samples were filled in 1.5 mm pathlength titanium double sector centerpieces (Nanolytics). All measurements were carried out using Rayleigh interference optics at 60,000 rpm rotor speed at 25 °C. The time evolution of sedimentation data was analyzed with SEDFIT software (version 15.01c)^48^. The weight concentration distribution of components, *c*(*s*_20,w_), was obtained as a function of the sedimentation coefficient. Here, the sedimentation coefficient was normalized to the value at 20 °C in pure water, *s*_20,w_.

### Analytical size-exclusion chromatography

Size exclusion chromatography to assess complex formation was performed using a Superdex 200 increase 10/300 (Cytiva). For sample preparation, 10 µM Kapβ2 and 0-50 µM MBP-PR18 were mixed at various molar ratios in a buffer containing 20 mM Hepes pH 7.4, 150 mM NaCl, 10% glycerol, 2 mM Mg(OAc)_2_, and 2 mM DTT. Thereafter, 500 μL of protein samples was loaded onto the column and eluted with buffer containing 20 mM Hepes pH 7.4, 150 mM NaCl, 10% glycerol, 2 mM Mg(OAc)_2_, and 2 mM DTT.

### NMR spectroscopy

NMR samples were prepared in 50 mM Tris buffer (pH 8.0), 150 mM NaCl. The protein concentration used was 190 µM. NMR experiments were performed on Bruker 600 MHz and 800 MHz NMR operated with Topspin software at 25 °C. The spectra were processed using the NMRPipe program^49^, and data analysis was performed using Sparky (Goddard TD, Kneller DG (1997) SPARKY 3, University of California, San Francisco. http://www.cgl.ucsf.edu/home/sparky). For resonance assignment, [^1^H-^15^N] HSQC, HNCO, HN(CO)CA, HNCA, CBCA(CO)NH, HNCACB, C(CO)NH, and HBHA(CO)NH spectra were recorded for ^13^C/^15^N-labeled protein.

### Fluorescence binding assay using Dnc-PR20

Kapβ2 Δloop or *Ce*Kapβ2 at various concentrations (10, 20, 30, 50, 100, 200, 300, 600, 1200, 1800 nM) was prepared in a buffer containing 20 mM HEPES buffer (pH 7.4), 150 mM NaCl, 2 mM MgCl_2_, 2 mM DTT, and 20% (w/v) glycerol with or without Dnc-PR20. The samples were measured in the fluorescence spectrum between 400 and 700 nm, excited by 340 nm light at 20 °C using an FP-8300 spectrofluorometer (JASCO Corp., Tokyo, Japan). Fluorescence spectra of the Kapβ2-DncPR20 interaction were obtained using the Spectra Manager software (JASCO Corp.) by subtracting the spectrum without Dnc-PR20 from the spectrum with Dnc-PR20. The change in fluorescence intensity at 550 nm depending on Kapβ2 concentration was analyzed as an index of the binding amount ratio using Microsoft Excel software (Microsoft).

## References

1. DeJesus-Hernandez, M. et al. Expanded GGGGCC Hexanucleotide Repeat in Noncoding Region of C9ORF72 Causes Chromosome 9p-Linked FTD and ALS. Neuron 72, 245–256 (2011).

2. Renton, A. E. et al. A hexanucleotide repeat expansion in C9ORF72 is the cause of chromosome 9p21-linked ALS-FTD. Neuron 72, 257–268 (2011).

3. Cleary, J. D. & Ranum, L. P. W. Repeat associated non-ATG (RAN) translation: New starts in microsatellite expansion disorders. Curr. Opin. Genet. Dev. 26, 6–15 (2014).

4. Zu, T. et al. Non-ATG-initiated translation directed by microsatellite expansions. Proc. Natl. Acad. Sci. U. S. A. 108, 260–265 (2011).

5. Mori, K. et al. The C9orf72 GGGGCC Repeat Is Translated into Aggregating Dipeptide-Repeat Proteins in FTLD/ALS. Science 339, 1335–1338 (2013).

6. Zhang, K. et al. The C9orf72 repeat expansion disrupts nucleocytoplasmic transport. Nature 525, 56–61 (2015).

7. Freibaum, B. D. et al. GGGGCC repeat expansion in C9orf72 compromises nucleocytoplasmic transport. Nature 525, 129–133 (2015).

8. Jovičić, A. et al. Modifiers of C9orf72 dipeptide repeat toxicity connect nucleocytoplasmic transport defects to FTD/ALS. Nat. Neurosci. 18, 1226–1229 (2015).

9. Shi, K. Y. et al. Toxic PRn poly-dipeptides encoded by the C9orf72 repeat expansion block nuclear import and export. Proc. Natl. Acad. Sci. U. S. A. 114, E1111–E1117 (2017).

10. Freibaum, B. D. & Taylor, J. P. The role of dipeptide repeats in C9ORF72-related ALS-FTD. Frontiers in Molecular Neuroscience vol. 10 at 10.3389/fnmol.2017.00035 (2017).

11. Cicardi, M. E. et al. The nuclear import receptor Kapβ2 modifies neurotoxicity mediated by poly(GR) in C9orf72-linked ALS/FTD. *Commun*. Biol. 7, 376 (2024).

12. Sato, T. et al. Axonal ligation induces transient redistribution of TDP-43 in brainstem motor neurons. Neuroscience 164, 1565–1578 (2009).

13. Springhower, C. E., Rosen, M. K. & Chook, Y. M. Karyopherins and condensates. Curr. Opin. Cell Biol. 64, 112–123 (2020).

14. Lee, B. J. et al. Rules for Nuclear Localization Sequence Recognition by Karyopherinβ2. Cell 126, 543–558 (2006).

15. Zhang, Z. C. & Chook, Y. M. Structural and energetic basis of ALS-causing mutations in the atypical proline-tyrosine nuclear localization signal of the Fused in Sarcoma protein (FUS). Proc. Natl. Acad. Sci. U. S. A. 109, 12017–12021 (2012).

16. Weis, K. Regulating access to the genome: Nucleocytoplasmic transport throughout the cell cycle. Cell 112, 441–451 (2003).

17. Görlich, D. & Kutay, U. Transport Between the Cell Nucleus and the Cytoplasm. Annu. Rev. Cell Dev. Biol. 15, 607–660 (1999).

18. Yoshizawa, T. et al. Nuclear Import Receptor Inhibits Phase Separation of FUS through Binding to Multiple Sites. Cell 173, 693–705.e22 (2018).

19. Nanaura, H. et al. C9orf72-derived arginine-rich poly-dipeptides impede phase modifiers. Nat. Commun. 1–12 (2021) doi:10.1038/s41467-021-25560-0.

20. Kwon, I. et al. Poly-dipeptides encoded by the C9orf72 repeats bind nucleoli, impede RNA biogenesis, and kill cells. Science 345, 1139–1145 (2014).

21. Mizielinska, S. et al. C9orf72 repeat expansions cause neurodegeneration in Drosophila through arginine-rich proteins. Science 345, 1192–1194 (2014).

22. Dormann, D. et al. ALS-associated fused in sarcoma (FUS) mutations disrupt Transportin-mediated nuclear import. EMBO J. 29, 2841–2857 (2010).

23. Gal, J. et al. Nuclear localization sequence of FUS and induction of stress granules by ALS mutants. Neurobiol. Aging 32, 2323.e27–2323.e40 (2011).

24. Ito, D., Seki, M., Tsunoda, Y., Uchiyama, H. & Suzuki, N. Nuclear transport impairment of amyotrophic lateral sclerosis-linked mutations in FUS/TLS. Ann. Neurol. 69, 152–162 (2011).

25. Cansizoglu, A. E., Lee, B. J., Zhang, Z. C., Fontoura, B. M. A. & Chook, Y. M. Structure-based design of a pathway-specific nuclear import inhibitor. Nat. Struct. Mol. Biol. 14, 452–454 (2007).

26. Wen, X., et al. Antisense Proline-Arginine RAN Dipeptides Linked to C9ORF72-ALS/FTD Form Toxic Nuclear Aggregates that Initiate In Vitro and In Vivo Neuronal Death. Neuron 84, 1213–1225 (2014).

27. Hao, Z. et al. Motor dysfunction and neurodegeneration in a C9orf72 mouse line expressing poly-PR. Nat. Commun. 10, (2019).

28. Tyzack, G. E. et al. Widespread FUS mislocalization is a molecular hallmark of amyotrophic lateral sclerosis. Brain 142, 2572–2580 (2019).

29. Tomita, S. Chemical tongues: biomimetic recognition using arrays of synthetic polymers. Polym. J. 54, 851–862 (2022).

30. Tomita, S. et al. Optical Fingerprints of Proteases and Their Inhibited Complexes Provided by Differential Cross-Reactivity of Fluorophore-Labeled Single-Stranded DNA. ACS Appl. Mater. Interfaces 11, 47428–47436 (2019).

31. Tomita, S. et al. Polymer-based chemical-nose systems for optical-pattern recognition of gut microbiota. Chem. Sci. (2022) doi:10.1039/d2sc00510g.

32. Li, Z., Askim, J. R. & Suslick, K. S. The Optoelectronic Nose: Colorimetric and Fluorometric Sensor Arrays. Chem. Rev. 119, 231–292 (2019).

33. Dam, J. & Schuck, P. Sedimentation velocity analysis of heterogeneous protein-protein interactions: Sedimentation coefficient distributions c(s) and asymptotic boundary profiles from Gilbert-Jenkins theory. Biophys. J. 89, 651–666 (2005).

34. Jafarinia, H., der Giessen, E. Van & Onck, P. R. Molecular basis of C9orf72 poly-PR interference with the β-karyopherin family of nuclear transport receptors. Sci. Rep. 12, 21324 (2022).

35. Miyagi, T. et al. Differential toxicity and localization of arginine-rich C9ORF72 dipeptide repeat proteins depend on de-clustering of positive charges. iScience 26, 106957 (2023).

36. Jafarinia, H., Van der Giessen, E. & Onck, P. R. C9orf72 polyPR directly binds to various nuclear transport components. bioRxiv (2023) doi:10.1101/2023.06.22.546068.

37. Xu, L. et al. C9orf72 poly(PR) aggregation in nucleus induces ALS/FTD-related neurodegeneration in cynomolgus monkeys. Neurobiol. Dis. 184, 106197 (2023).

38. Hutten, S. et al. Nuclear Import Receptors Directly Bind to Arginine-Rich Dipeptide Repeat Proteins and Suppress Their Pathological Interactions. Cell Rep. 33, (2020).

39. Bourgeois, B. et al. Nonclassical nuclear localization signals mediate nuclear import of CIRBP. Proc. Natl. Acad. Sci. 117, 8503–8514 (2020).

40. Mannen, T., Yamashita, S., Tomita, K., Goshima, N. & Hirose, T. The Sam68 nuclear body is composed of two RNasesensitive substructures joined by the adaptor HNR NPL. J. Cell Biol. 214, 45–59 (2016).

41. Kato, M., Lin, Y. & McKnight, S. L. Cross-β polymerization and hydrogel formation by low-complexity sequence proteins. Methods 126, 3–11 (2017).

42. Baek, Y. S. & Park, Y. K. Intensity-based holographic imaging via space-domain Kramers– Kronig relations. Nat. Photonics 15, 354–360 (2021).

43. Lee, M., Kunzi, M., Neurohr, G., Lee, S. S. & Park, Y. Hybrid machine-learning framework for volumetric segmentation and quantification of vacuoles in individual yeast cells using holotomography. Biomed. Opt. Express 14, 4567–4578 (2023).

44. Schindelin, J., et al. Fiji: An open-source platform for biological-image analysis. Nat. Methods 9, 676–682 (2012).

45. Axelrod, D., Koppel, D. E., Schlessinger, J., Elson, E. & Webb, W. W. Mobility measurement by analysis of fluorescence photobleaching recovery kinetics. Biophys. J. 16, 1055–1069 (1976).

46. Nehls, S. et al. Dynamics and retention of misfolded proteins in native ER membranes. Nat. Cell Biol. 2, 288–295 (2000).

47. Kemmer, G. & Keller, S. Nonlinear least-squares data fitting in Excel spreadsheets. Nat. Protoc. 5, 267–281 (2010).

48. Schuck, P. Size-distribution analysis of macromolecules by sedimentation velocity ultracentrifugation and Lamm equation modeling. Biophys. J. 78, 1606–1619 (2000).

49. Delaglio, F. et al. NMRPipe: A multidimensional spectral processing system based on UNIX pipes. J. Biomol. NMR 6, 277–293 (1995).

