## Extended data fig. 1-15 and table 1 for "Conserved loop of a phase modifier endows protein condensates with fluidity"

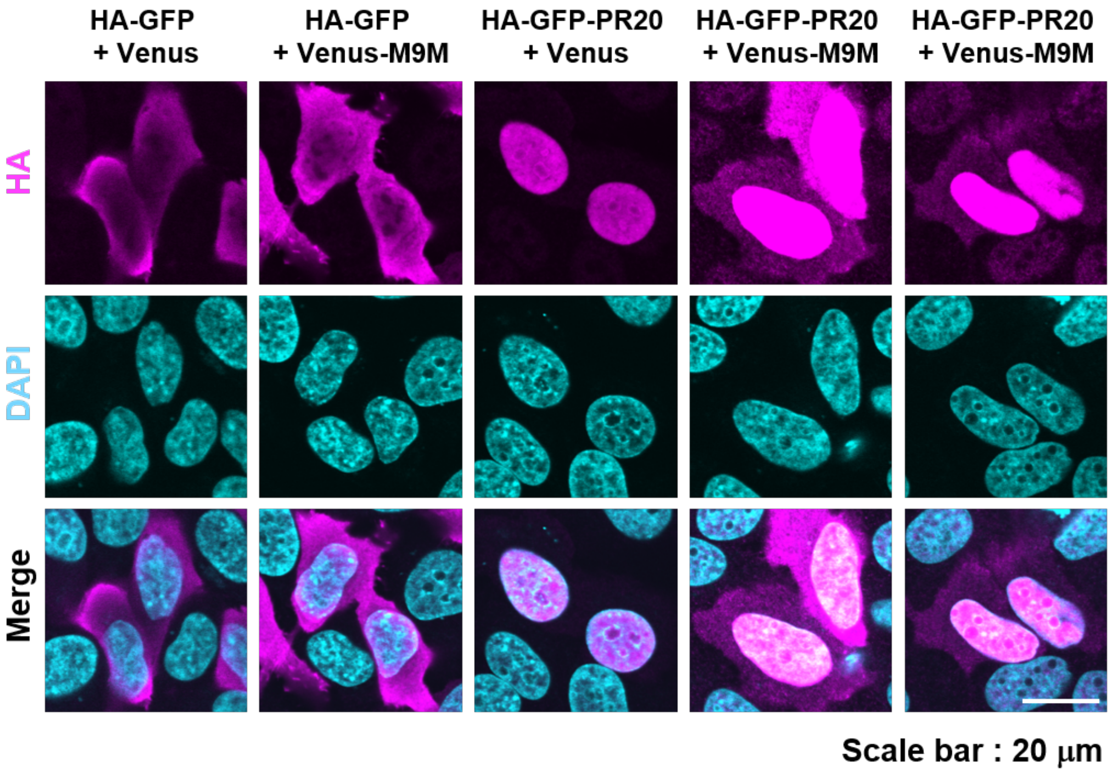


**Extended Data Fig. 1: The subcellular localization of PR20 expressed in HeLa cell in the presence of Kapβ2 specific inhibitor.**

Confocal immunofluorescence images showing the subcellular localization of HA-tagged GFP or GFP-PR20 (in purple) in HeLa cells. Cells were costained with DAPI (in blue) to highlight the nucleus. Kapβ2 specific inhibitor, M9M, was coexpressed as Venus fusion protein (+ Venus-M9M).

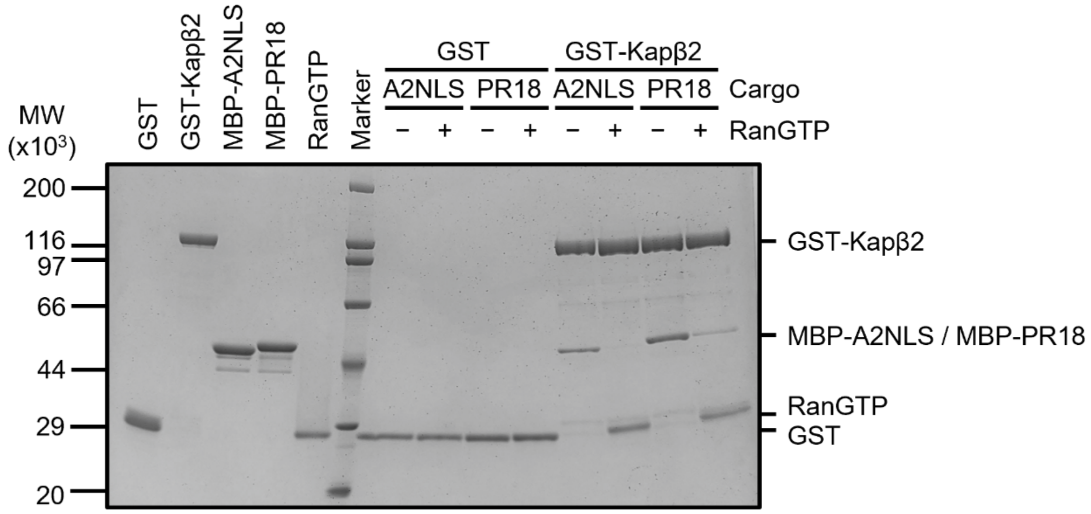


**Extended Data Fig. 2: Kapβ2 cargo unloading by RanGTP in vitro.**

Image of the entire gel showing the pull-down binding assay of MBP-hnRNPA2NLS or MBP-PR18 with immobilized GST-Kapβ2 in the presence of RanGTP. Right portion of this gel is shown as the Figure 1B.

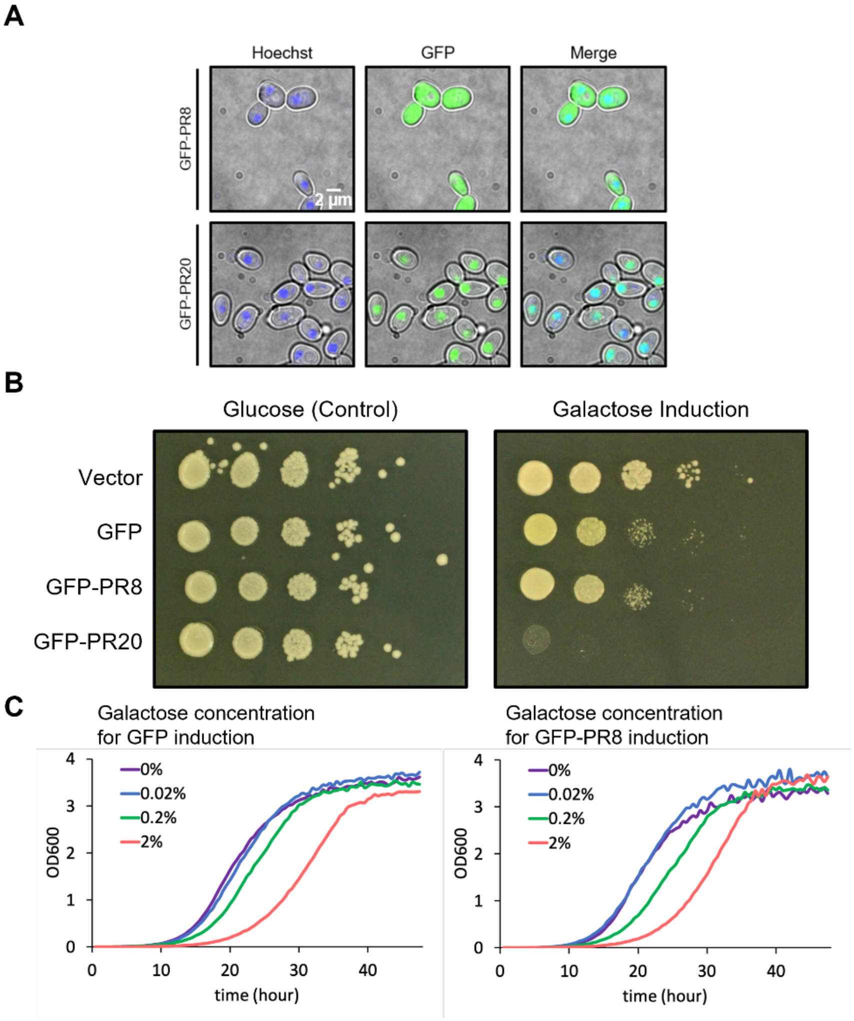


**Extended Data Fig. 3: Evaluation of subcellular localization of PR poly-dipeptides and growth defect by PR poly-dipeptide expression in yeast.**

**A**. Fluorescence images showing the subcellular localization of GFP-PR8 and GFP-PR20 (in green) in living yeast cells.

**B**. Spot assay to examine cell growth when GFP, GFP-PR8, or GFP-PR20 was expressed in yeast cells. Each gene is under *GAL1* promoter control. The concentrations of serial dilution spots were from left to right: $10^7$, $10^6$, $10^5$, $10^4$, and $10^3$ cells/mL. The concentration of glucose and galactose in the agar medium was 2%.

**C.** Dose-dependent effect of PR poly-dipeptide expression on yeast growth. The percentage numbers indicate galactose concentration for induction of GFP (left) or GFP-PR8 (right) expression.

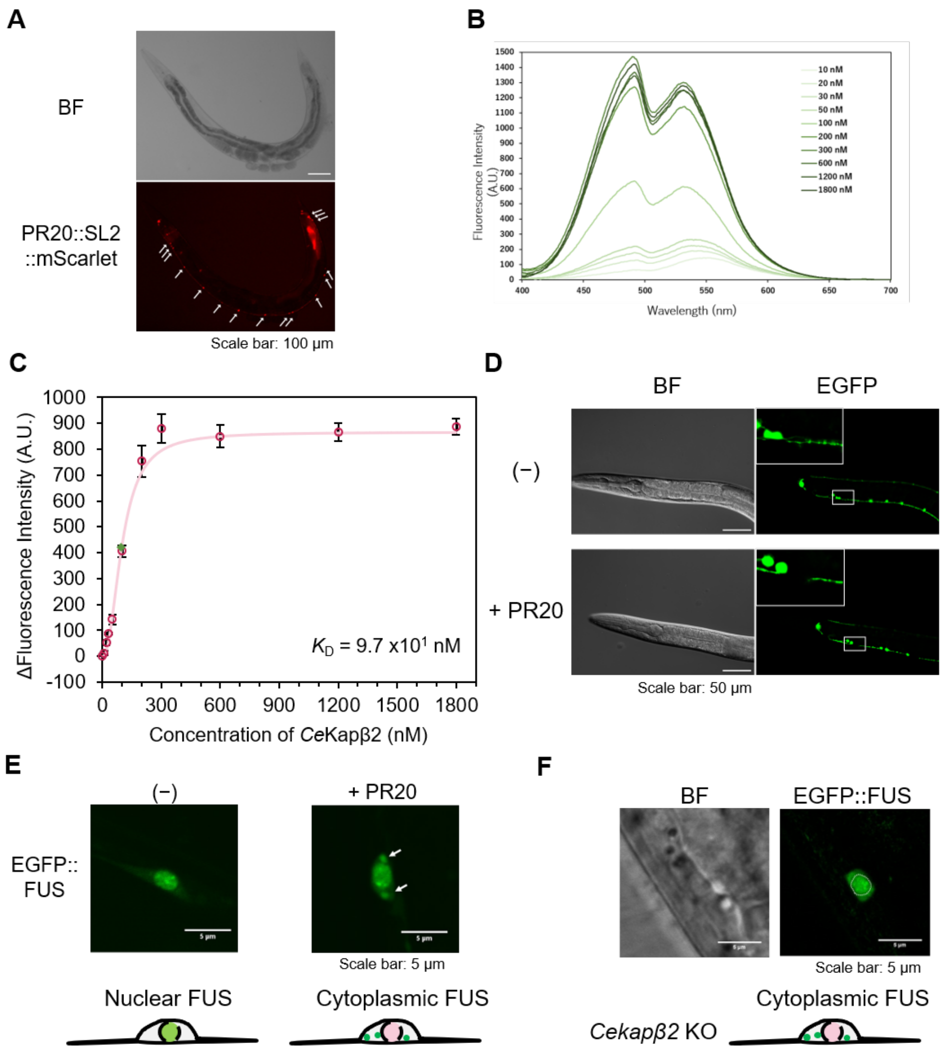


**Extended Data Fig. 4: The effect of PR20 expression in *C. elegans*.**

**A**. PR20 expression in motor neurons of *C. elegans*. The expression of PR20 was visualized by PR20::SL2::mScarlet gene under control of *unc-47* promoter at adult stage. Arrows indicate the site of expression. Scale bar, 100 µm.

**B**. Fluorescence emission spectra of Dnc-PR20 in the presence of *Ce*Kapβ2. PR20 (300 nM) modified with Dnc group (Dnc-PR20) was mixed with 10–1800 nM *Ce*Kapβ2 and Dnc group was excited with a 340 nm light.

**C**. The in vitro interaction analysis between *Ce*Kapβ2 and Dnc-PR20. Dnc-PR20 was titrated with *Ce*Kapβ2 and the change in fluorescence intensity at 550 nm was measured. The concentration with half maximum fluorescence intensity ($K_D$) was determined from the fitting curve and is indicated by the green triangle. Data are indicated as mean ± SD from two independent experiments.

**D**. Neuronal degeneration by PR20 expression. Motor neurons were visualized by the expression of GFP by *unc-25* promoter. Missing neurites are shown in the inset. Scale bar, 50 µm.

**E**. Abnormal localization of FUS by PR20 expression. EGFP::FUS was expressed in motor neurons by *unc-47* promoter. Abnormal localization of FUS condensation in cytoplasm is indicated by arrows. Scale bar, 5 μm.

**F**. Abnormal localization of FUS by deletion of *Cekapβ2*. EGFP::FUS was uniformly spread throughout the cytoplasm in addition to the nucleus. Dotted circle indicates the nucleus. Scale bar, 5 μm.

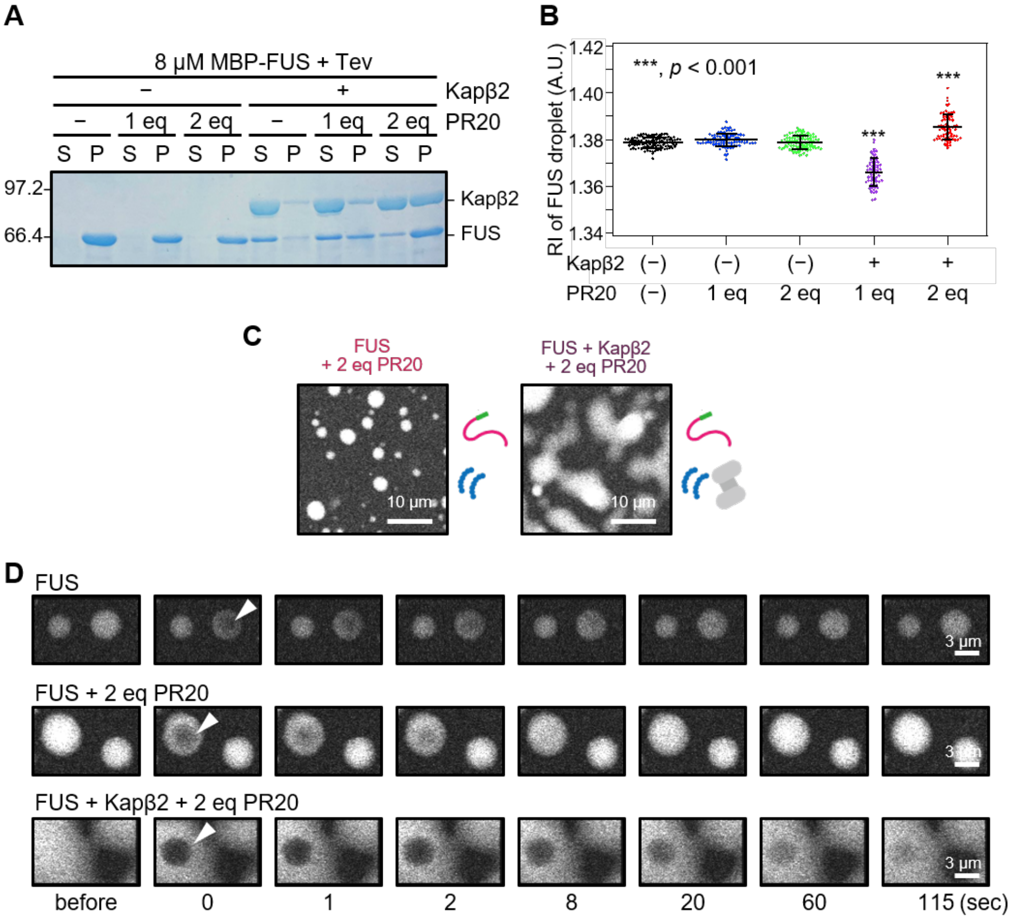


**Extended Data Fig. 5: Effect of excess PR20 to FUS droplet in the presence of Kapβ2 .**

**A**. Proteins in supernatant and precipitant of centrifuged FUS droplets were confirmed by SDS-PAGE. FUS droplet solutions were prepared as in Fig. 1E. After incubation, the droplet solutions were separated into supernatant (S) as a soluble fraction and precipitant (P) as an insoluble fraction by centrifugation, and the fractions were analyzed by SDS-PAGE.

**B**. Quantitative analysis of refractive index images of FUS droplets at different concentrations of PR20 peptide in the absence and presence of Kapβ2. Data are mean±SD. MBP-FUS, $n$ = 113 droplets from five independent experiments; MBP-FUS + 1 eq PR20, $n$ = 100 droplets from four independent experiments; MBP-FUS + 2 eq PR20, $n$ = 101 droplets from four independent experiments; MBP-FUS + Kapβ2 + 1 eq PR20, $n$ = 66 droplets from five independent experiments; MBP-FUS + Kapβ2 + 2 eq PR20, $n$ = 74 droplets from six independent experiments.

**C**. Confocal fluorescent images of FUS droplet in the presence of both PR20 and Kapβ2. Droplet formation was triggered by addition of 4% (w/v) PEG8000 to FUS solutions containing 9 μM MBP-FUS, 1 μM MBP-FUS-ATTO488, and 20 μM PR20 in the absence or presence of 10 μM Kapβ2. Fluorescent signal excited with a 473 nm laser. Scale bar, 10 μm.

**D**. Confocal fluorescent images of FUS droplet before and after photobleaching related to Fig. 1G.

Photobleaching was performed in the area indicated by the white arrowheads on the droplets 30 minutes after the droplet formation, and images were taken over time. FUS droplet without Kapβ2 and PR20 (upper panels), FUS droplet with 20 μM PR20 (middle panels), FUS droplet with 10 μM Kapβ2 and 20 μM PR20 (lower panels), respectively. Scale bar, 3 μm.

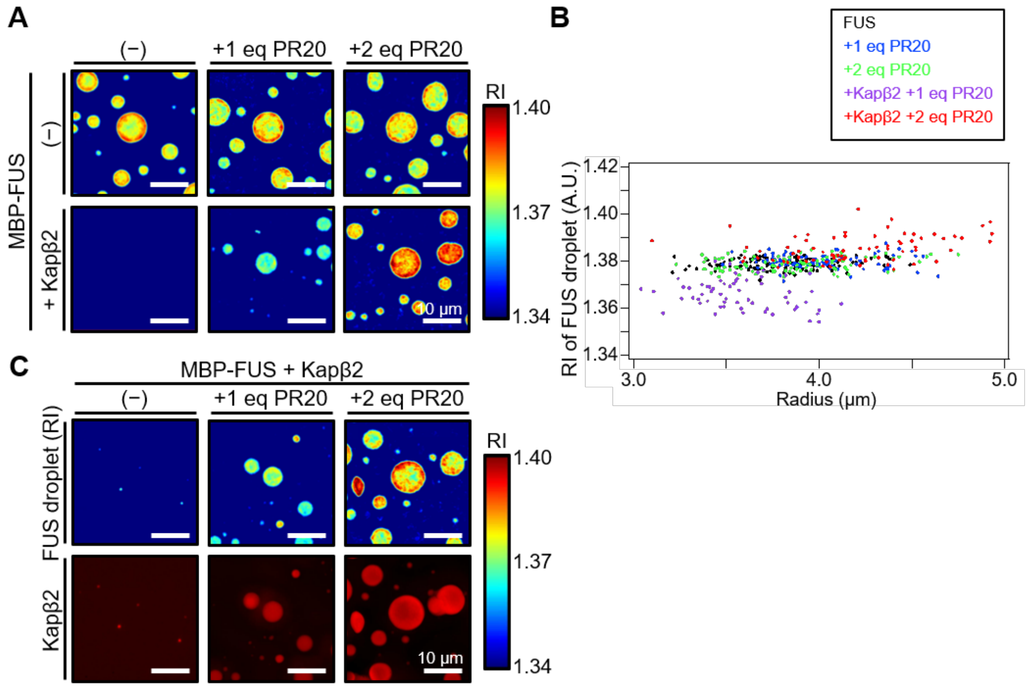


**Extended Data Fig. 6: Optical diffraction tomography analysis of FUS droplet in the presence of PR20 and Kapβ2.**

**A**. ODT images of FUS droplets in the presence of varying concentrations of PR20 in the absence and presence of Kapβ2. Zoom out of the view in Fig. 1F. Scale bar, 10 μm.

**B**. The relationship between droplet radius and its refractive index. Radius vs RI plots of datapoints in Extended Data Fig. 5B. Although particle size and RI are weakly correlated under some conditions, the RI differences induced by PR20 are much more significant than size dependency in this analysis. MBP-FUS, $n$ = 113 droplets from five independent experiments; MBP-FUS + 1 eq PR20, $n$ = 100 droplets from four independent experiments; MBP-FUS + 2 eq PR20, $n$ = 101 droplets from four independent experiments; MBP-FUS + Kapβ2 + 1 eq PR20, $n$ = 66 droplets from five independent experiments; MBP-FUS + Kapβ2 + 2 eq PR20, $n$ = 74 droplets from six independent experiments.

**C**. ODT and fluorescence imaging of FUS droplets with or without PR20 and Kapβ2. Zoom out of the view in Fig. 1H. Scale bar, 10 μm.

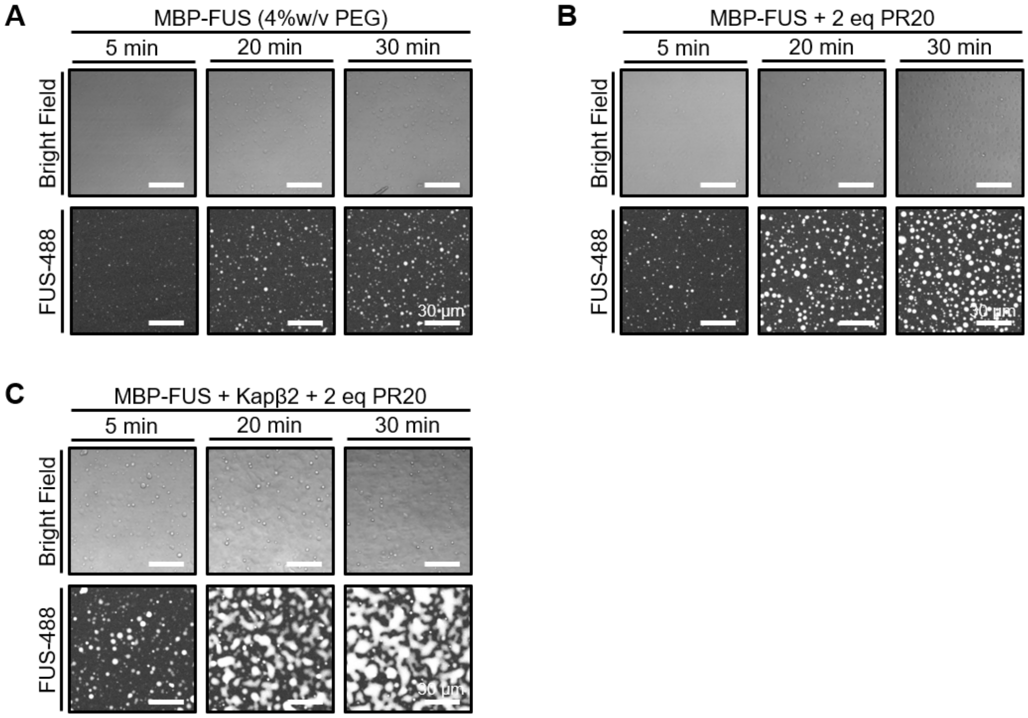


**Extended Data Fig. 7: FUS droplet formation and aging.**

**A**. Confocal fluorescent images of FUS droplet. Droplet formation was triggered by addition of 4%(w/v) PEG8000 to FUS solutions containing 9 μM MBP-FUS and 1 μM MBP-FUS-ATTO488. Bright Field, differential interference contrast image; FUS-488, fluorescent signal excited with a 473 nm laser. Scale bars: 30 μm.

**B**. Confocal fluorescent images of FUS droplet with 20 μM PR20.

**C**. Confocal fluorescent images of FUS droplet with 10 μM Kapβ2 and 20 μM PR20.

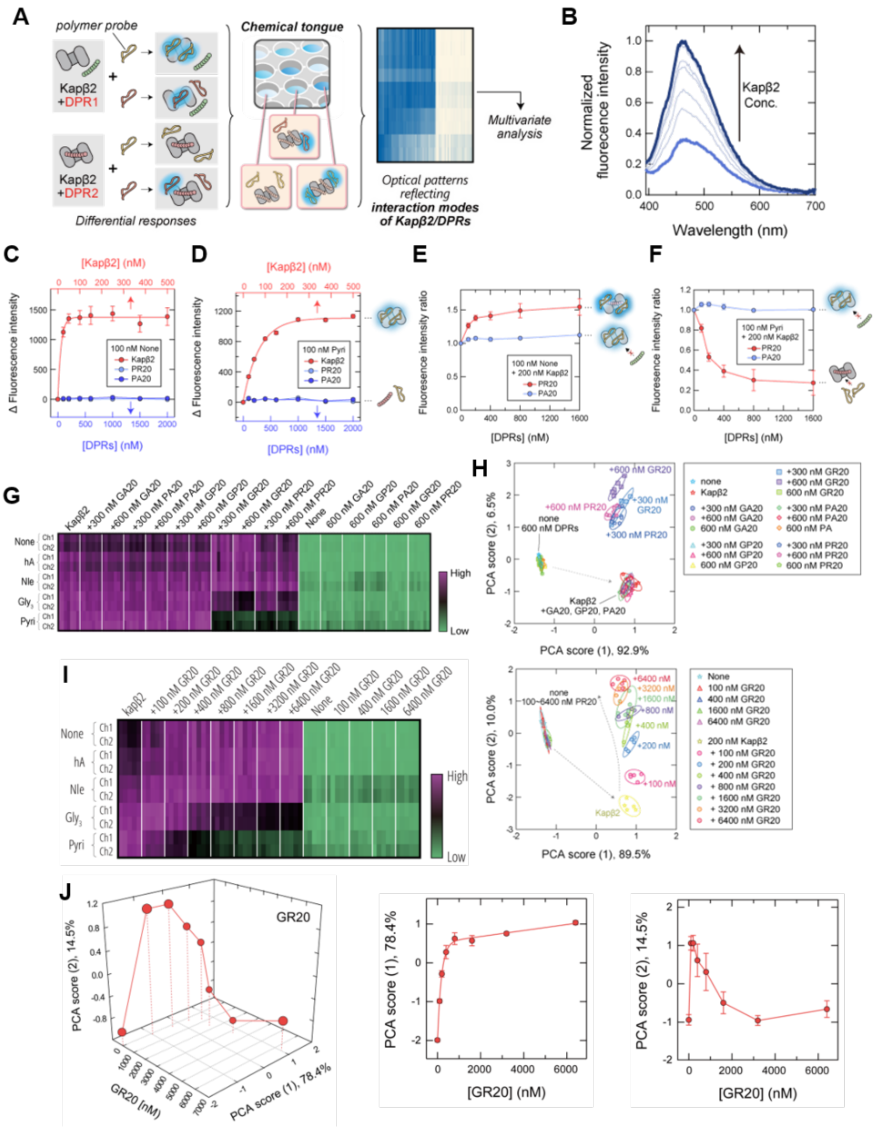

**Extended Data Fig. 8: Chemical tongue analysis on Kapβ2 and DPRs.**

**A**. Schematic illustration of the chemical tongue strategy to extract interaction information between Kapβ2 and DPRs. Upon binding with Kapβ2, the polymer probes with aggregation-induced emission dyes increase in fluorescence intensity, but remain inert to DPRs, thus, the formation of Kapβ2/DPRs complexes modulates these intensities by covering Kapβ2's interaction sites, and the resultant optical patterns reflect the interaction modes between Kapβ2 and DPRs.

**B**. Normalized fluorescence spectra of TPE-None (100 nM) upon addition of Kapβ2 (0-500 nM).

**C**. Binding isotherms of TPE-None (100 nM) upon addition of Kapβ2 (red), PR20 (light blue), PA20 (blue).

**D**. Binding isotherms of TPE-Pyri (100 nM) upon addition of Kapβ2 (red), PR20 (light blue), PA20 (blue).

**E**. The fluorescence intensity ratios of TPE-None (100 nM) and Kapβ2 (200 nM) upon addition of

DPRs; PR20 (red) and PA20 (light blue).

**F**. The fluorescence intensity ratios of TPE-Pyri (100 nM) and Kapβ2 (200 nM) upon addition of DPRs; PR20 (red) and PA20 (light blue).

**G**. Heat map of the fluorescence response patterns of Kapβ2 in the absence and presence of GA20, PA20, GP20, GR20 and PR20.

**H**. PCA score plot of Kapβ2 in the absence and presence of GA20, PA20, GP20, GR20 and PR20.

**I**. Heat map of the fluorescence response patterns and PCA score plot of Kapβ2 in the absence and presence of GR20.

**J**. PCA scores for Kapβ2 in the absence and presence of GR peptide plotted against GR concentration.

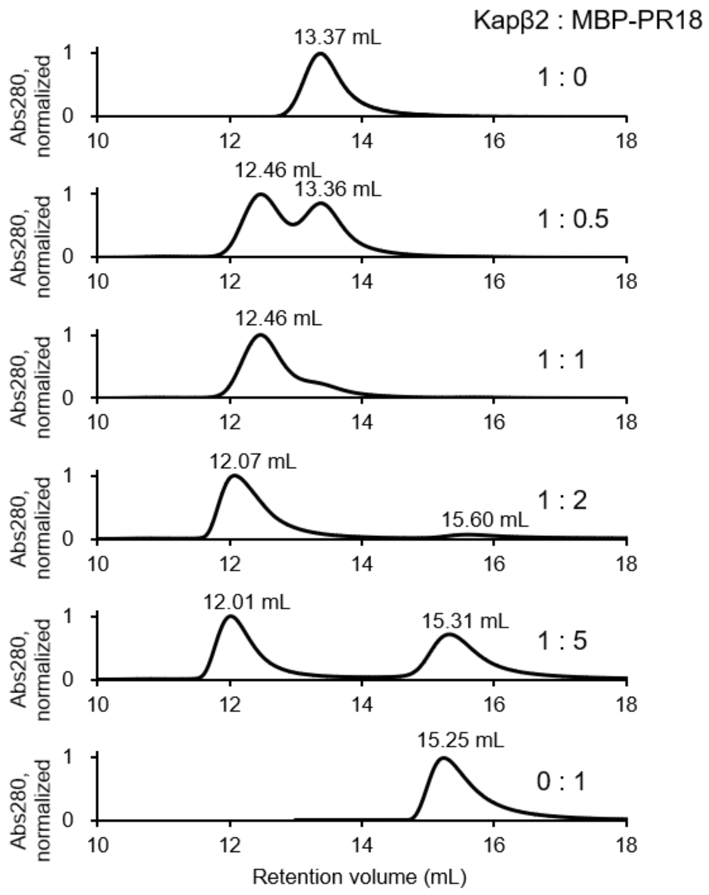


**Extended Data Fig. 9: Analytical size-exclusion chromatography for Kapβ2 and MBP-PR18.** Kapβ2 and MBP-PR18 were mixed at the molar ratios shown in right and run on size-exclusion chromatography. The absorbance of the eluate was normalized to the maximum value. Elution volumes of peaks were labeled.

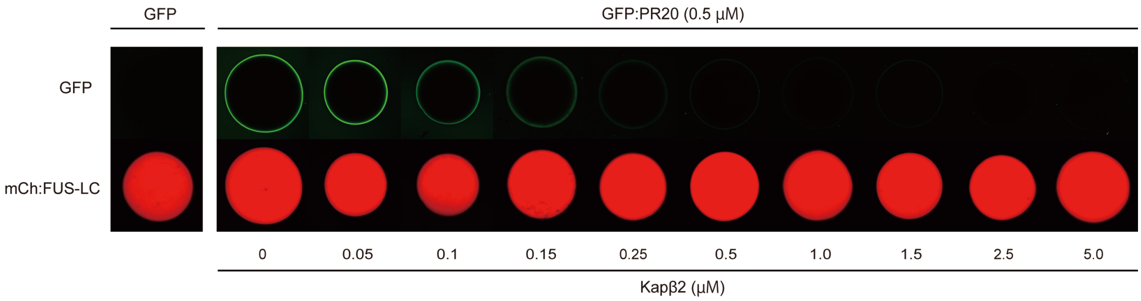


**Extended Data Fig. 10: Hydrogel binding assay for FUS-LC in the absence and presence of Kapβ2 and PR20.**

GFP:PR20 were applied to mCh:FUS-LC hydrogel droplets in the absence and presence of Kapβ2. Hydrogel droplets of mCh:FUS-LC (lower images) were incubated with 0.5 µM of GFP (left panel) or 0.5 µM of GFP:PR20 (right panel) in the presence of different concentrations of Kapβ2 (left to right: from 0 to 5.0 µM).

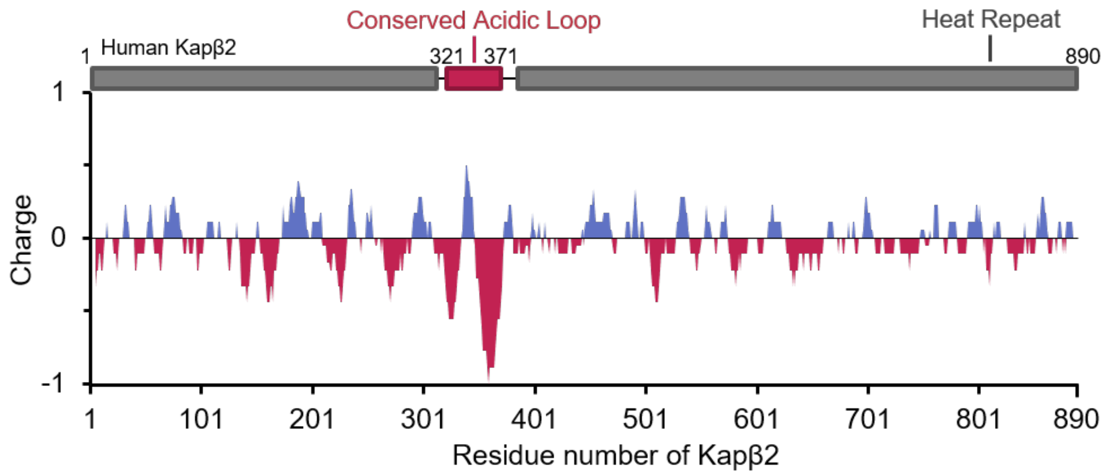


**Extended Data Fig. 11: Charge distribution on the primary sequence of Kapβ2.**

Surface charge was predicted based on amino acid sequence of Kapβ2 using EMBOSS: charge server (window length 9; https://www.bioinformatics.nl/cgi-bin/emboss/charge?_pref_hide_optional=0). The negative value means predicted negative charge of the region. Acidic loop region (321−371) in schematic representation of Kapβ2 domain composition is highlighted in red.

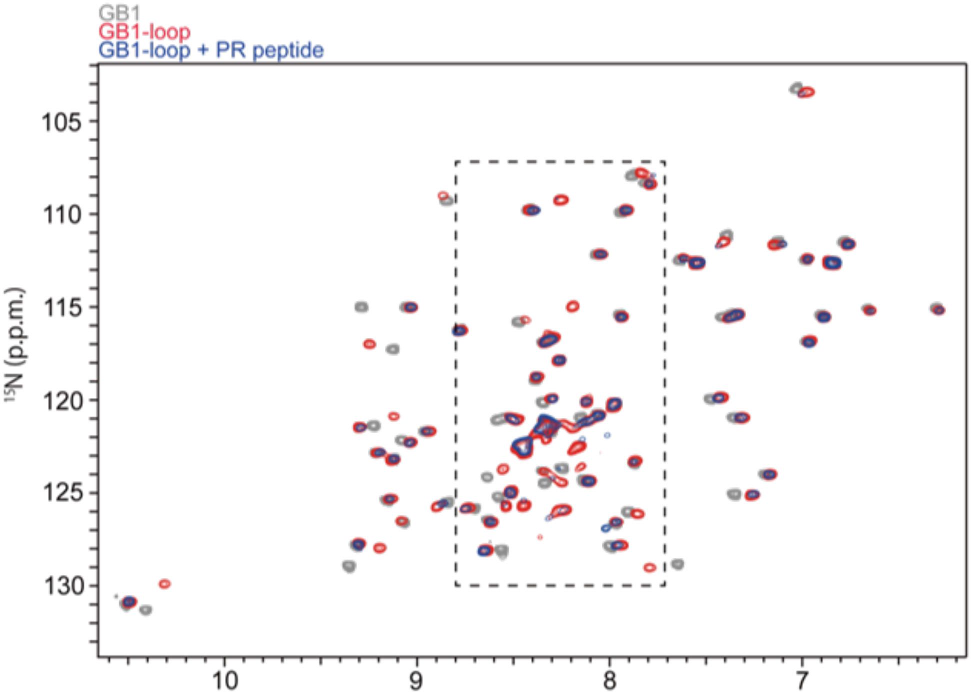


**Extended Data Fig. 12: Entire $^{1}$H-$^{15}$N HSQC spectra of GB1 and GB1-loop in the absence and presence of PR20.**

The data correspond to the selected region of the spectra shown in Fig. 3C.

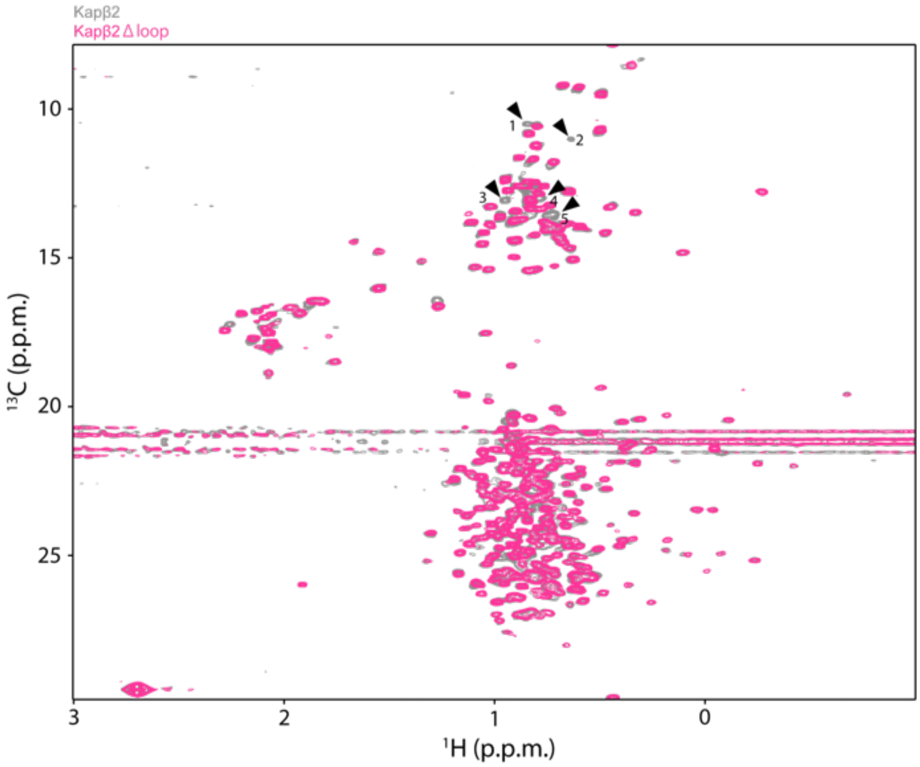


**Extended Data Fig. 13: $^{1}$H-$^{13}$C HMQC spectra of Kapβ2 WT and Δloop.**

$^{1}$H-$^{13}$C-correlated methyl NMR spectra of [U-$^{2}$H; Ile-δ1-$^{13}CH_3$; Leu, Val-$^{13}CH_3$/$^{12}CH_3$]-labeled Kapβ2 WT (gray) and Kapβ2 Δloop (pink). In the spectrum of Kapβ2 Δloop, several resonances presumably from methyl groups in the loop disappear, while majority of the resonances appear at the positions corresponding to the WT resonances, indicating that the core structure is preserved in the Kapβ2 Δloop.

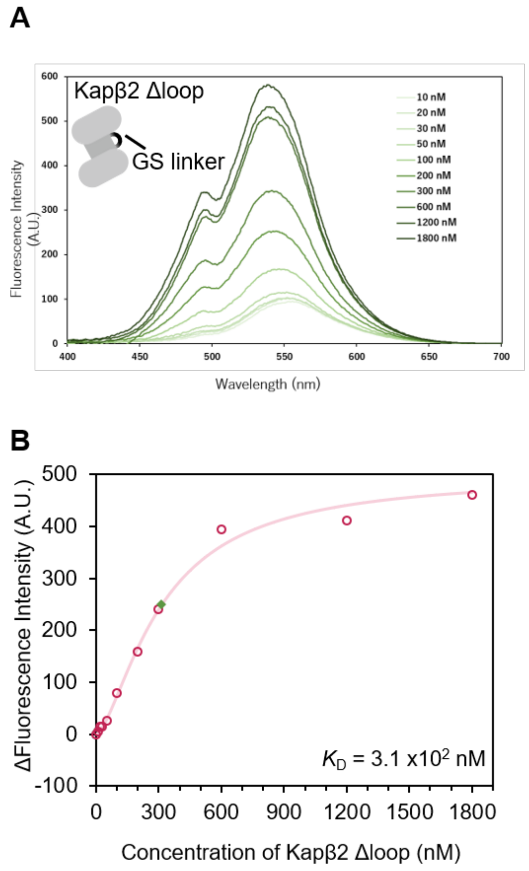


**Extended Data Fig. 14: Binding analysis between PR20 and Kapβ2 using fluorescence.**

**A**. Fluorescence emission spectra of Dnc-PR20 in the presence of Kapβ2. Dnc-PR20 (300 nM) was mixed with 10–1800 nM Kapβ2 and Dnc group was excited with a 340 nm light.

**B**. The in vitro interaction analysis between Kapβ2 and Dnc-PR20 peptide. Dnc-PR20 was titrated with Kapβ2 and the change in fluorescence intensity at 550 nm was measured. The concentration with half maximum fluorescence intensity ($K_D$) was determined from the fitting curve and is indicated by the green diamond.

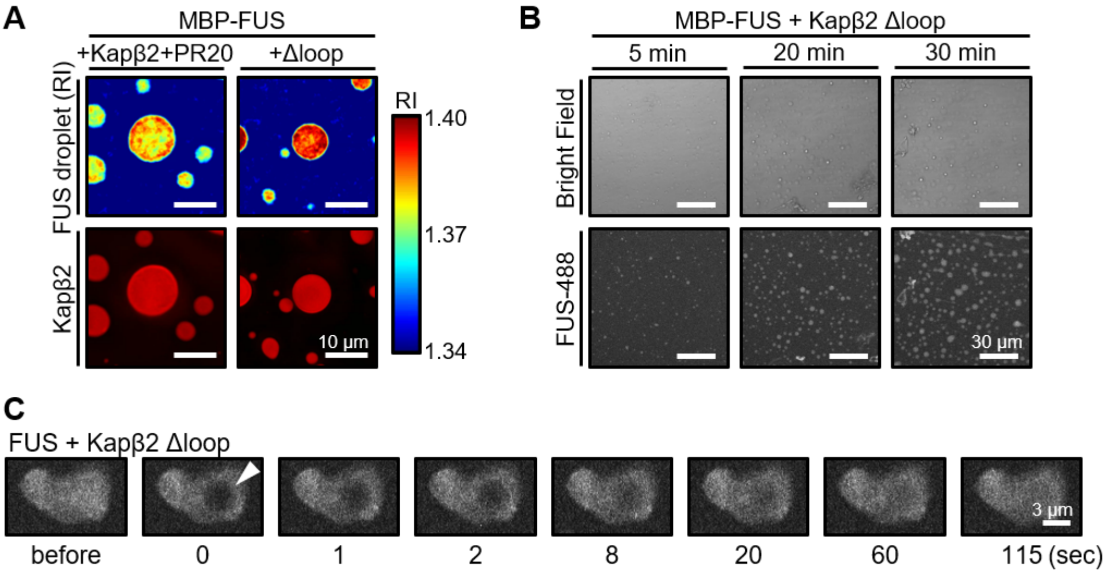


**Extended Data Fig. 15: Effect of Δloop mutation on Kapβ2 to the function as a phase modifier.**

**A**. ODT and fluorescence images of FUS droplets in the presence of Kapβ2 and 2 eq PR20 or Kapβ2 Δloop. Zoom out of the RI image view in Fig. 4A. Scale bar, 10 µm.

**B**. Confocal fluorescent images of FUS droplet with 10 µM Kapβ2 Δloop. Scale bar, 30 µm.

**C**. Confocal fluorescent images of FUS droplet with 10 µM Kapβ2 Δloop before and after photobleaching related to Fig. 4C. Photobleaching was performed in the area indicated by the white arrowheads on the droplet 30 minutes after the droplet formation, and images were taken over time. Scale bar, 3 µm.

**Extended Data Table. 1: Statistics of counted droplets.**

| | average RI | STDEV | $r$/μm (Ave) | STDEV | N |
|---|---|---|---|---|---|
| MBP-FUS | 1.3787 | 0.0022 | 3.7985 | 0.2607 | 113 |
| MBP-FUS + 1 eq PR20 | 1.3798 | 0.0027 | 4.0004 | 0.2494 | 100 |
| MBP-FUS + 2 eq PR20 | 1.3787 | 0.0028 | 3.8616 | 0.3128 | 101 |
| MBP-FUS + Kapβ2 + 1 eq PR20 | 1.3661 | 0.0059 | 3.4959 | 0.3670 | 66 |
| MBP-FUS + Kapβ2 + 2 eq PR20 | 1.3853 | 0.0055 | 4.2064 | 0.3788 | 74 |
| MBP-FUS + Kapβ2 Δloop | 1.3891 | 0.0039 | 3.9951 | 0.3047 | 59 |

The average RI value and particle size of each droplet were measured based on the RI images related to Fig. 1F, 4B, Extended Data Fig. 5B, and 6B.
